# An intestinal Sir2-HSF1-ATGL1 pathway regulates lipolysis in *C. elegans*

**DOI:** 10.1101/2024.04.11.588856

**Authors:** Milán Somogyvári, Saba Khatatneh, Gábor Hajdú, Bar Sotil, József Murányi, Csaba Sőti

## Abstract

Proteostasis maintenance and lipid metabolism are critical for survival and promote longevity, however, their coordination is largely unclear. Here we show that the heat shock factor HSF-1 and the proteostasis state regulates lipolysis in *C. elegans*. We find that in response to starvation, the sirtuin 1 ortholog SIR-2.1 activates lipolysis by upregulation of the adipose triglyceride lipase ATGL-1. In feeding worms, intestinal HSF-1 represses ATGL-1 expression and lipolysis via the microRNA system. In starving worms, SIR-2.1 suspends a *miR-53-* mediated suppression of lipolysis by inhibiting its HSF-1-dependent expression. The apparent antagonism of SIR-2.1 and HSF-1, distinct from their synergism at heat shock promoters suggests a context-specific regulation of HSF-1 by SIR-2.1. We demonstrate that the SIR-2.1 and protein kinase A pathways are both indispensable, and independently converge on ATGL- 1 for lipolysis. HSF-1 activation by proteostasis disturbances inhibits starvation-induced lipid mobilization, whereas its age-related decline limits fat deposition through *atgl-1*. Our findings reveal a crosstalk between proteostasis and lipid/energy metabolism, which may modulate stress resilience and aging.

## Introduction

Lipids are diverse constituents crucial to maintain the health for all organisms. They are involved in signal transduction, membrane structure and fluidity, as well as energy storage and mobilization^1^. Fat energy stores, mainly as triacylglycerol, serve as a primary source of energy during periods of fasting, starvation or caloric restriction. During these, the body alters its metabolism to mobilize and break down stored triglycerides (TG) into free fatty acids (FFAs) through the process of lipolysis, which can then be used as fuel by various tissues using β-oxidation^2^. Caloric restriction and its mimetics, as well as regulated fasting can extend lifespan and improve metabolic parameters, including lipid profiles ^3–5^. Additionally, intermittent fasting and calorie restriction may lead to enhanced cardiovascular health by positively influencing lipid markers, which may reduce the risk of cardiovascular diseases^6^. Furthermore, it has been found that lipids are involved in genetic pathways related to lifespan regulation^7,8^.

Lipolysis is a complex and highly regulated process in the roundworm *C. elegans*, involving a network of signaling pathways and several lipases. Although *C. elegans* does not have a dedicated fat storage tissue, its intestine and hypodermis possess all the conserved lipid metabolism pathways and regulators^9^. Notably, several lines of research in worms uncovered that longevity pathways, fat metabolism and lifespan determination are interconnected^10–14^. The conserved adipose triglyceride lipase (ATGL), the key enzyme of lipolysis possesses a crucial role in metabolic homeostasis and exhibits intricate transcriptional, post- transcriptional and post-translational regulation involving FOXO, sirtuins, insulin signaling, protein kinase A (PKA) and the AMPK-activated protein kinase (AMPK)^2,15^. In *C. elegans*, ATGL-1 is the major lipase that mediates starvation-induced lipolysis, which involves its phosphorylation by PKA^16^. PKA also regulates fat mobilization during cold adaptation *via* the upregulation of hormone sensitive lipase ortholog *hosl-1*^17^. Moreover, ATGL-1 is critical for survival during the dauer stage through the AMPK pathway^18,19^ and mediates longevity in response to dietary restriction and reduced insulin-like signaling ^11^. Although these findings underscore the importance of ATGL-1 in nematodal energy homeostasis, its regulation is not entirely clear.

The sirtuin 1 (SIRT1) NAD^+^-dependent deacetylase is conserved from yeast to mammals. It is critically involved in various cellular processes such as gene expression, metabolism^20,21^, and longevity^22,23^. For instance, overexpression of SIRT1 in transgenic mice causes lifespan extension and delays the onset of age-related diseases^24^. Moreover, SIRT1 plays diverse beneficial roles in metabolic homeostasis and protects from the harmful effects of obesity and insulin resistance^25^. SIRT1 regulates adipocyte lipolysis by controlling the FOXO1-dependent transcription of ATGL^26^. The *C. elegans* ortholog SIR-2.1 has been found to regulate stress response and aging in a context-dependent fashion^27,28^. Part of its effect is due to its interaction with the FOXO ortholog DAF-16 transcription factor^29^. SIRT1 activates the heat shock transcription factor HSF1 in mammals^30^ and SIR-2.1 was demonstrated to synergize with its activity in *C. elegans*^31^. Moreover, SIR-2.1 was implicated in the fasting-induced inhibition of SREBP-mediated lipid synthesis ^32^. However, the impact of SIR-2.1 on lipolysis is unknown, yet.

HSF1 plays a crucial role in guarding the proteome and in the cellular response to proteotoxic stress by regulating the expression of multiple heat-shock proteins (HSPs)^33,34^. *C. elegans* HSF-1 was found to regulate global stress-responsive gene expression to promote stress-resistance and survival^35^ and housekeeping functions in development, metabolism, and longevity^36,37^. Dietary deprivation was shown to extend lifespan and to protect against proteotoxicity in an HSF-1-dependent manner^38^. HSF-1 also regulates longevity through nutrient signaling pathways, like the *C. elegans* insulin/IGF-1 and target of rapamycin (TOR) signaling^39,40^. While our work was in progress, a recent study reported that loss of *C. elegans hsf-1* indirectly impairs nutrient absorption, which in turn activates the intracellular lipid surveillance pathway and restoration of fat stores via NHR-49^41^. However, it is unknown whether HSF-1 might directly regulate lipid mobilization.

In this study we introduce a novel lipolysis-regulatory axis in the worm intestine, where HSF-1 negatively regulates lipid mobilization by suppressing *atgl-1* expression through the microRNA system. In response to starvation, SIR-2.1 activates lipolysis by suspending the HSF-1-mediated inhibition, whereas intestinal proteotoxic stress impairs lipid mobilization via HSF-1. Our results reveal a previously unknown crosstalk between lipid and protein homeostasis.

## Results

### SIR-2.1 upregulates ATGL-1 expression to activate lipolysis during starvation

In order to investigate the previously reported involvement of SIR-2.1 in lipid storage during starvation^32^, wildtype, *sir-2.1* knockout, and *sir-2.1* overexpressing animals fed by either empty vector (EV) or *sir-2.1(RNAi)* from L4 larval stage for 2 days were then subjected to starvation for 18 hours. Lipid content was evaluated by Oil Red O (ORO) staining. Our results confirmed that the lack of *sir-2.1* activity either by mutation or by gene silencing impaired the reduction of fat stores in response to starvation (Figure 1A and B). Moreover, SIR-2.1 overexpression further reduced lipid stores in both fed and starved animals. Evaluation of free fatty acid (FFA) content showed that both starvation and SIR-2.1 overexpression in fed nematodes elevated FFA levels, which were completely inhibited by *sir-2.1(RNAi)* (Figure 1C). The elevated FFA levels in the absence of feeding are likely due to increased triglyceride breakdown, which suggests a regulatory role for SIR-2.1. in lipid mobilization.

**Figure 1.**
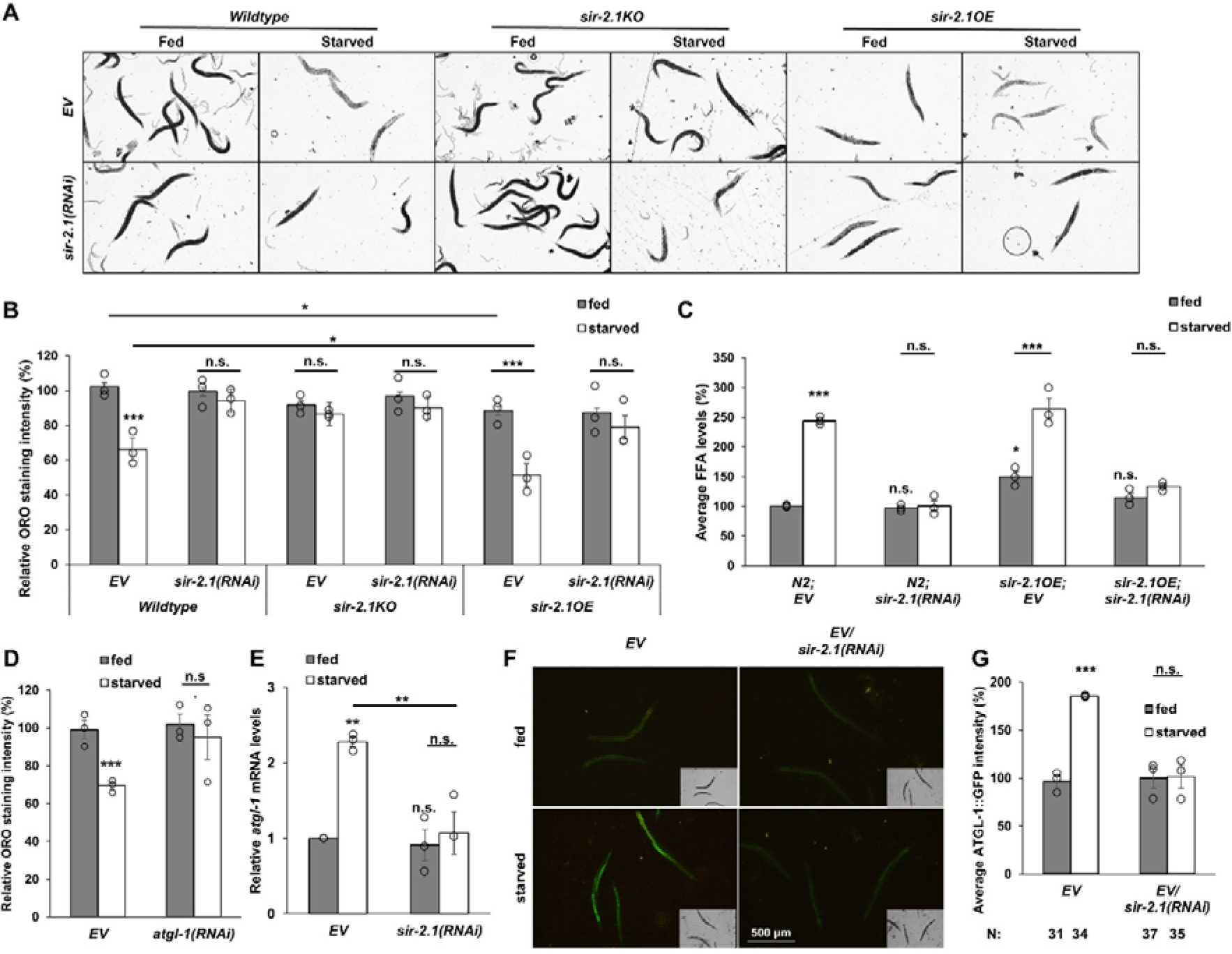
SIR-2.1 activation mediates starvation-induced lipolysis through *atgl-1* mRNA upregulation. (A-B) Representative micrographs (A) and statistical evaluation (B) showing the effect of *sir-2.1* RNAi, mutation and overexpression on starvation-induced lipid mobilization by ORO staining (C) Levels of free fatty acids (FFAs) in response to starvation were measured in animals either overexpressing *sir-2.1* and/or having it silenced by RNAi. (D) Evaluation of the ORO staining of animals treated by empty vector (EV) or *atgl-1(RNAi)*. (E) Relative mRNA levels of *atgl-1* on EV and *sir-2.1(RNAi)*. (F-G) Induction of the *atgl- 1p::atgl-1::gfp* transgene by starvation and the effect of *sir-2.1* silencing. Data are expressed as mean ± SEM of three independent experiments. Circles represent the mean values of individual experiments. N, number of animals measured/condition. p values were obtained by two-way ANOVA using the Fisher’s LSD test. n.s.: not significant; *: p<0.05; **: p<0.01; ***: p<0.001.

To test a potential *sir-2.1*-dependent regulation, we explored the effector lipase(s) responsible for lipolysis. The hormone sensitive lipase *hosl-1*^17^, the enoyl-CoA hydratase *ech-4*^42^, and the triglyceride lipase *lips-6*^43^ have all been reported previously to affect lipid content and/or mobilization. RNAi against these enzymes did not affect ORO staining upon starvation (Figure S1A). The adipose triglyceride lipase *atgl-1* however, proved to be necessary for starvation-induced lipid mobilization, and *atgl-1* silencing phenocopied the loss of *sir-2.1* activity (Figure 1D). We then asked whether starvation and *sir-2.1* regulates *atgl-1* expression by measuring *atgl-1* mRNA levels and GFP fluorescence intensity in an ATGL-1::GFP reporter strain. These experiments show that both *atgl-1* mRNA and protein levels are upregulated by starvation in a *sir-2.1*-dependent manner (Figure 1E-G). Moreover, *sir-2.1* was also required for the starvation-induced upregulation of *lips-6* and *hosl-1* mRNAs (Figure S1B, C), indicating its critical role in efficient mobilization of fat stores.

### Loss of HSF-1 function is required for SIR-2.1-dependent lipolysis in the intestine

Our observation on the *sir-2.1*-dependent upregulation of *atgl-1* and other mRNAs led us to investigate the possible involvement of various transcription factors reported to play a role in lipid metabolism. These were the nuclear hormone receptor *nhr-49* (promoting beta oxidation ^44^), the helix-loop-helix-type transcriptional factor *hlh-30* (inducing the expression of lipases, like *lipl-1*^45^) and the FOXA ortholog *pha-4* (required for lipase expressions^46^. RNAi against these transcription factors did not affect lipid content of fed and starved nematodes (Figure S2A). Additionally, we also investigated the possible involvement of stress-responsive transcription factors in *sir-2.1*-dependent lipolysis. Knock-out mutation of the FOXO transcription factor DAF-16 had no effect on the starvation-induced lipid mobilization in EV and *sir-2.1*-fed worms (Figure S2B). Likewise, mutation or silencing of the heat-shock transcription factor HSF-1 *per se*, did not affect the lipid content of fed and starved worms. However, it restored the ability of *sir-2.1* mutant and RNAi-fed animals to mobilize lipid resources in response to starvation (Figure 2A). This result was confirmed by measurements of the free fatty acid content of identically treated animals (Figure 2B). Interestingly, *hsf- 1(RNAi)* not only restored lipolysis, but appeared to phenocopy the increased FFA content observed in the SIR-2.1 overexpressing strain (cf. Figures 2B and 1C). To test if ATGL-1 is indeed the effector in the mechanism thus outlined, we combined RNAi against *atgl-1* and *hsf-1*. Here, *hsf-1(RNAi)* failed to restore lipid mobilization, confirming *atgl-1*’s role as the downstream effector of this pathway (Figure S2C).

**Figure 2.**
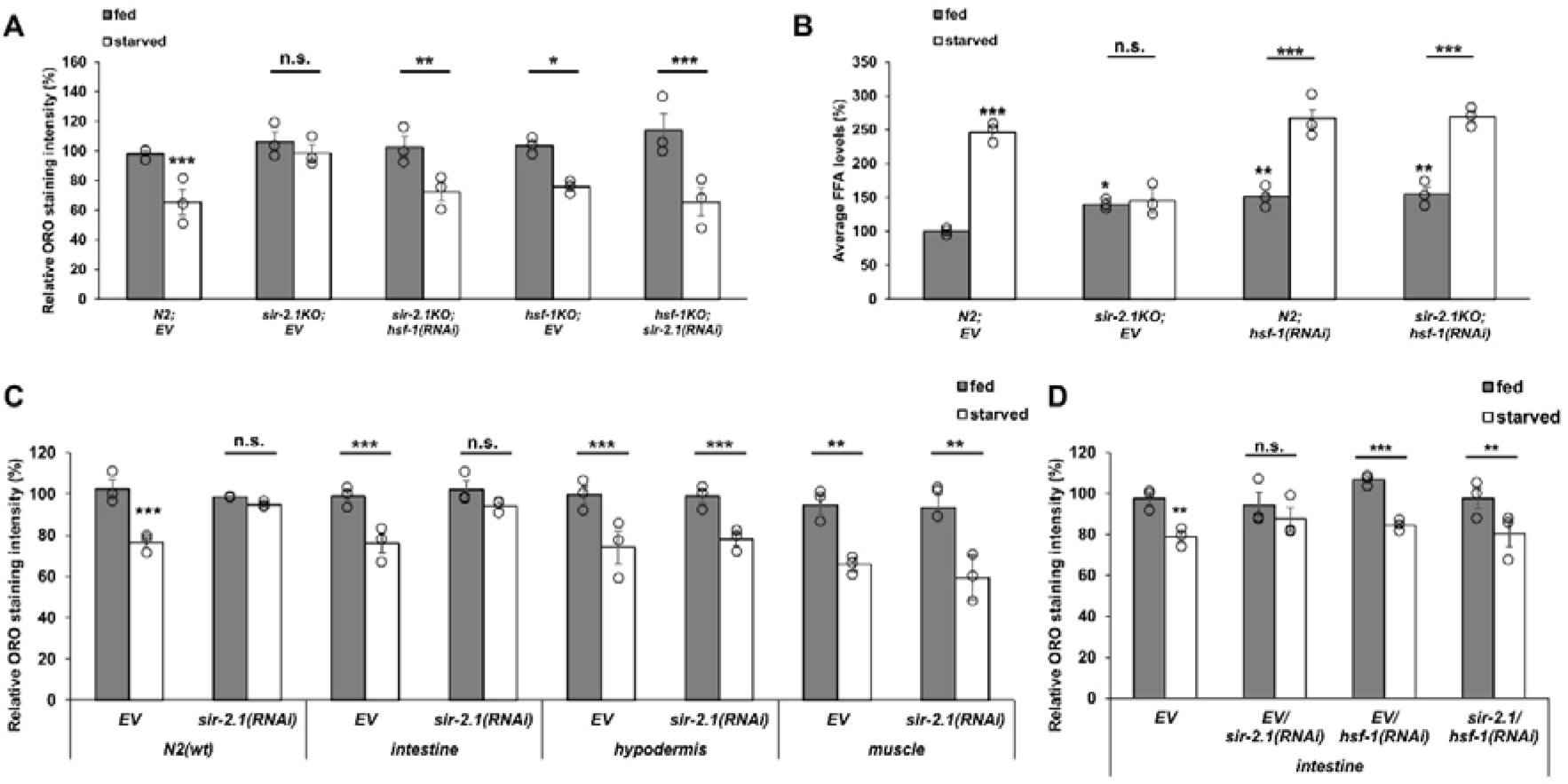
Loss of HSF-1 function is required for SIR-2.1-dependent lipolysis in the intestine. (A) Epistasis analysis of loss of *hsf-1* and *sir-2.1* in starvation-induced lipid mobilization. (B) Quantification of free fatty acid levels in response to starvation upon *hsf-1* and/or *sir-2.1(RNAi)*. (C) The effect of *sir-2.1(RNAi)* on starvation-induced lipid mobilization in wildtype and in different tissue-specific RNAi strains. (D) Lipid mobilization upon *sir-2.1* and/or *hsf-1(RNAi)* treatment in the intestinally RNAi-sensitive strain. Data are expressed as mean ± SEM of three independent experiments. Circles represent the mean values of individual experiments. p values were obtained by two-way ANOVA using the Fisher’s LSD test. n.s.: not significant; *: p<0.05; **: p<0.01; ***: p<0.001.

In order to identify the tissue(s) where the *sir-2.1/hsf-1/atgl-1* pathway regulates lipolysis, we utilized several strains with tissue-selective sensitivity for RNAi in the intestine^47^, the hypodermis and body wall muscle^48^. We found that RNAi against *sir-2.1* only inhibited lipolysis in the intestine (Figure 2C). Furthermore, *hsf-1(RNAi)* restored lipolysis in intestine- specific *sir-2.1*-silenced worms, indicating a cell-autonomous interaction of SIR-2.1 and HSF- 1 in the gut (Figure 2D).

### HSF-1 represses ATGL-1 expression and lipolysis through the microRNA system

Results of the epistasis analysis suggested that downstream from *sir-2.1*, the inhibition of *hsf-1* activity is necessary for efficient lipolysis upon starvation. We reasoned that in this scenario, HSF-1 activation might prevent starvation-induced lipolysis. First, we took use of the increased genetic dosage of HSF-1 in two distinct HSF-1 overexpressor strains. We observed a complete inhibition of starvation-induced lipolysis in both strains (Fig 3A). At non-stress conditions, a conserved interaction with the HSP-90 chaperone complex keeps HSF-1 under inhibition^49,50^. Therefore, we activated HSF-1 by *hsp-90* RNAi knockdown and found an absence of lipid mobilization upon starvation, which was restored in *hsf-1* mutants (Fig 3B). Silencing *hsp-90* was also sufficient to impair starvation-induced expression of the ATGL-1::GFP reporter strain (Fig 3C). Thus, these data establish that HSF-1 activity prevents lipolysis through ATGL-1, which effect is in turn counteracted by SIR-2.1 upon starvation.

**Figure 3.**
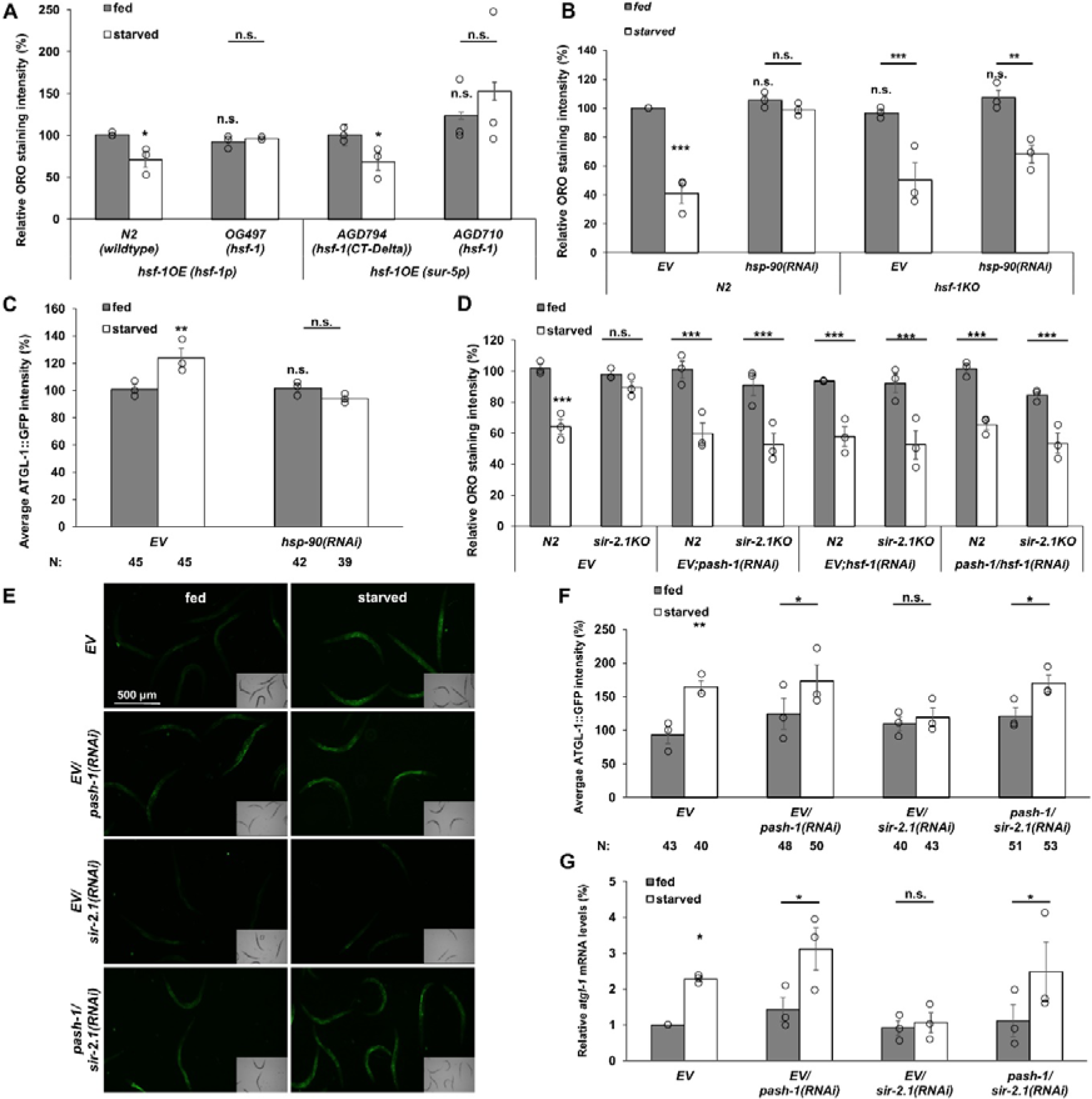
HSF-1 inhibits lipid mobilization and ATGL-1 expression through the microRNA system. (A) Increased gene dosage of HSF-1 prevents lipid mobilization in response to starvation. *hsf-1* transgene driven by its own promoter is compared to N2 wildtype control, whereas *hsf-1* driven by a *sur-5* promoter is compared to its own control expressing the truncated *hsf-1(CT-Delta)*. (B) Activation of HSF-1 by *hsp-90* knockdown prevents lipid mobilization in response to starvation. (C) The effect of *hsp-90* silencing on the expression of the *atgl-1p::atgl-1::gfp* transgene (D) The effect of silencing microRNA maturation gene *pash-1* on lipid mobilization in wildtype and *sir-2.1* knockout animals. Representative fluorescence microscopy image (E) and quantification of (F) the effect of *pash-1* and/or *sir-2.1* silencing on the expression of the *atgl-1p::atgl-1::gfp* transgene. (G) Effect of *pash-1* and/or *sir-2.1* silencing on starvation-induced *atgl-1* mRNA levels. Data are expressed as mean ± SEM of three independent experiments. Circles represent the mean values of individual experiments. N, number of animals measured/condition. p values were obtained by two-way ANOVA using the Fisher’s LSD test. n.s.: not significant; *: p<0.05; **: p<0.01; ***: p<0.001.

Next, we aimed to identify the mechanism by which HSF-1 inhibits lipolysis. A genome-wide study identified a plethora of HSF-1 target genes but *atgl-1* is not on the list ^36^, suggesting an indirect regulation. Importantly, HSF-1 is also responsible for transcriptional activation of various microRNAs ^35^. Since we detected an inhibition of *atgl-1* mRNA expression upon *sir- 2.1(RNAi)* (Figure 1E), we considered the microRNA system as a mediator of HSF-1’s effect on lipid mobilization. To test this hypothesis, we utilized RNAi against key components of the *C. elegans* miRNA maturation process. RNAi against the Drosha ortholog *pash-1*, involved in the maturation of pri-miRNAs into pre-miRNAs, resulted in a complete phenocopy of *hsf-1* mutation or silencing by similarly restoring lipid mobilization in *sir-2.1* knock-out mutants (Figure 3D). This result was confirmed by *dcr-1(RNAi)*, inhibiting the next step of miRNA maturation (Figure S3A). We further investigated the role of the microRNA system by utilizing the strain expressing the ATGL-1::GFP fusion protein. We found that blocking microRNA maturation *per se* seemed to non-significantly induce GFP signals in fed animals, but it restored ATGL-1:GFP induction upon starvation in the presence of *sir- 2.1(RNAi)* (Figure 3E and F). Likewise, silencing *pash-1* restored the *sir-2.1* RNAi-dependent inhibition of *atgl-1* and *lips-6* mRNA induction by starvation (Figures 3G and S3B). Finally, *pash-1* RNAi restored the inhibition of lipolysis by HSF-1 overexpression (Figure S3C, D) or by *hsp-90* silencing (Figure S3E). These data demonstrate a role for the microRNA system in the HSF-1-dependent regulation of starvation-induced lipid mobilization process.

Among the HSF-1-regulated microRNAs, some were suggested to participate in the regulation of several important life processes, like intracellular signaling, post-embryonic development, cell death, aging and importantly, fatty acid metabolism ^35^. Using the TargetScanWorm^51^ (targetscan.org) miRNA binding prediction algorithm we identified one particular microRNA, namely *mir-53*, to bind to the 3’-UTR of the *atgl-1* mRNA. From its pre-miRNA, two distinct miRNA transcripts are spliced, namely the 24-bp *F36H1.8a* and the 22-bp *F36H1.8b*, only the former was identified to bind to the *atgl-1* 3’UTR (Figure 4A). Other microRNAs were also reported to be involved in lipid metabolism, like *mir-60*, and *mir-75*^52^. Of the latter two microRNAs, *mir-75 T24D8.8b* splice variant was also found to be a promising candidate for silencing *atgl-1* expression due to its predicted ability to bind to its 3’-UTR (Figure 4A). Hence, we investigated *sir-2.1*-dependent lipolysis in the respective miRNA mutants. The resulting data show, that the lack of *miR-53* completely, whereas that of *miR-60* and *miR-75* partially restored lipid mobilization in *sir-2.1-*silenced starved nematodes (Figure 4B). Further studies using GFP-fusion promoter reporter strains revealed, that from the three, only *mir-53* promoter activity was reduced by starvation in a *sir-2.1*- and *hsf-1*- dependent manner, while *mir-75* promoter activity was maintained by HSF-1 and dropped in *sir-2.1(RNAi)*-treated starved worms to a similar level to that of *hsf-1(RNAi)*-treated animals (Figure 4C, D). The induction of the *miR-60* promoter was unaffected by the treatments (Figure 4E). Due to its complete lipolysis inhibitory effect and responsiveness to starvation, we focussed on the regulation of *miR-53* by SIR-2.1 and HSF-1 in subsequent experiments.

**Figure 4.**
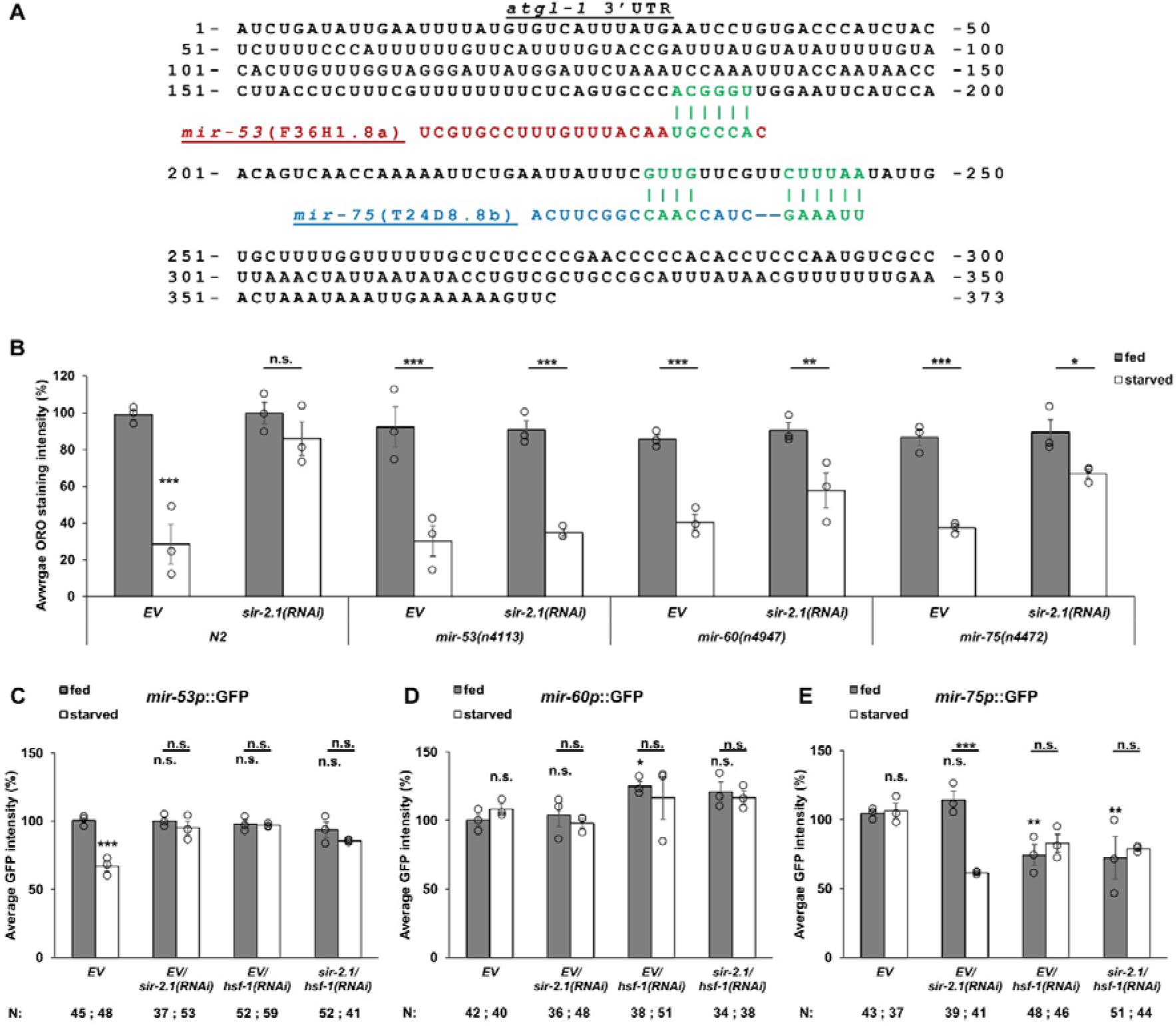
Inhibition of *miR-53* mediates starvation-induced lipolysis. (A) The sequence of *atgl-1*’s 3’UTR with predicted binding sites for *mir-53 F36H1.8a* and *mir-75 T24D8.8b*. (B) Lipid mobilization in response to starvation in different microRNA mutants. (C-E) Average GFP intensities of *mir-53* (C), *mir-60* (D) and *mir-75* (E) promoter reporter strains in response to starvation upon *sir-2.1* and/or *hsf-1* silencing. Data are expressed as mean ± SEM of three independent experiments. Circles represent the mean values of individual experiments. N, number of animals measured/condition. p values were obtained by two-way ANOVA using the Fisher’s LSD test. n.s.: not significant; *: p<0.05; **: p<0.01; ***: p<0.001.

### A context-dependent interaction of SIR-2.1 and HSF-1 regulates *mir-53* expression during starvation

Mammalian SIRT1 activates HSF1 upon heat stress by deacetylating a specific lysine residue^30^, which leads to a prolonged binding to target heat shock gene promoters, such as Hsp70. Moreover, heat and caloric restriction synergizes to upregulate the mRNA of Hsp70 orthologs in *C. elegans*, which requires *sir-2.1*^31^. Our findings suggested that during starvation, SIR-2.1 may liberate an HSF-1-dependent inhibition of lipolysis. To gain an insight into the unusual interaction of SIR-2.1 and HSF-1, we examined the promoter activities of *hsp-70 (C12C8.1)* and *mir-53* in response to heat shock and starvation, respectively, in the same experiment. Consistent with the previous study^31^ we found that both heat shock and starvation activated the *hsp-70* promoter, and both required *sir-2.1* and *hsf-1* (Figure 5A and B). However, only starvation, but not heat shock reduced the activation of the *mir-53* promoter, which was dependent on both *sir-2.1* and *hsf-1* (Figure 5C, D, note that though the experimental setup is the same, experiments presented in Fig. 5D are different from those of Fig. 4C). Contrary to our expectations, the knockdown of *hsf-1* was not sufficient to restore the activity of *mir-53* promoter *per se* or in *sir-2.1(RNAi)*-fed nematodes, which suggests a more complex regulation. Therefore, we aimed to assess *miR-53* RNA expression in *sir-2.1-* and *hsf-1*-silenced worms under starvation. Due to the short sequence of the mature *F36H1.8a* microRNA, RT-qPCR was used to determine the larger, 99-bp *mir-53* pre-miRNA (*F36H1.8*) levels. We observed an *hsf-1*-dependent upregulation of the transcript in starved *sir-2.1*- silenced nematodes (Figure 5E). Although the exact mechanism is unknown, it confirms that both the *mir-53* promoter as well as the stability/expression of its pre-RNA appears to be regulated. Altogether, these results suggest that during starvation, SIR-2.1 activation induces the HSF-1-dependent upregulation of *hsp-70* mRNA expression, while hindering that of the *pre-miR-53* RNA.

**Figure 5.**
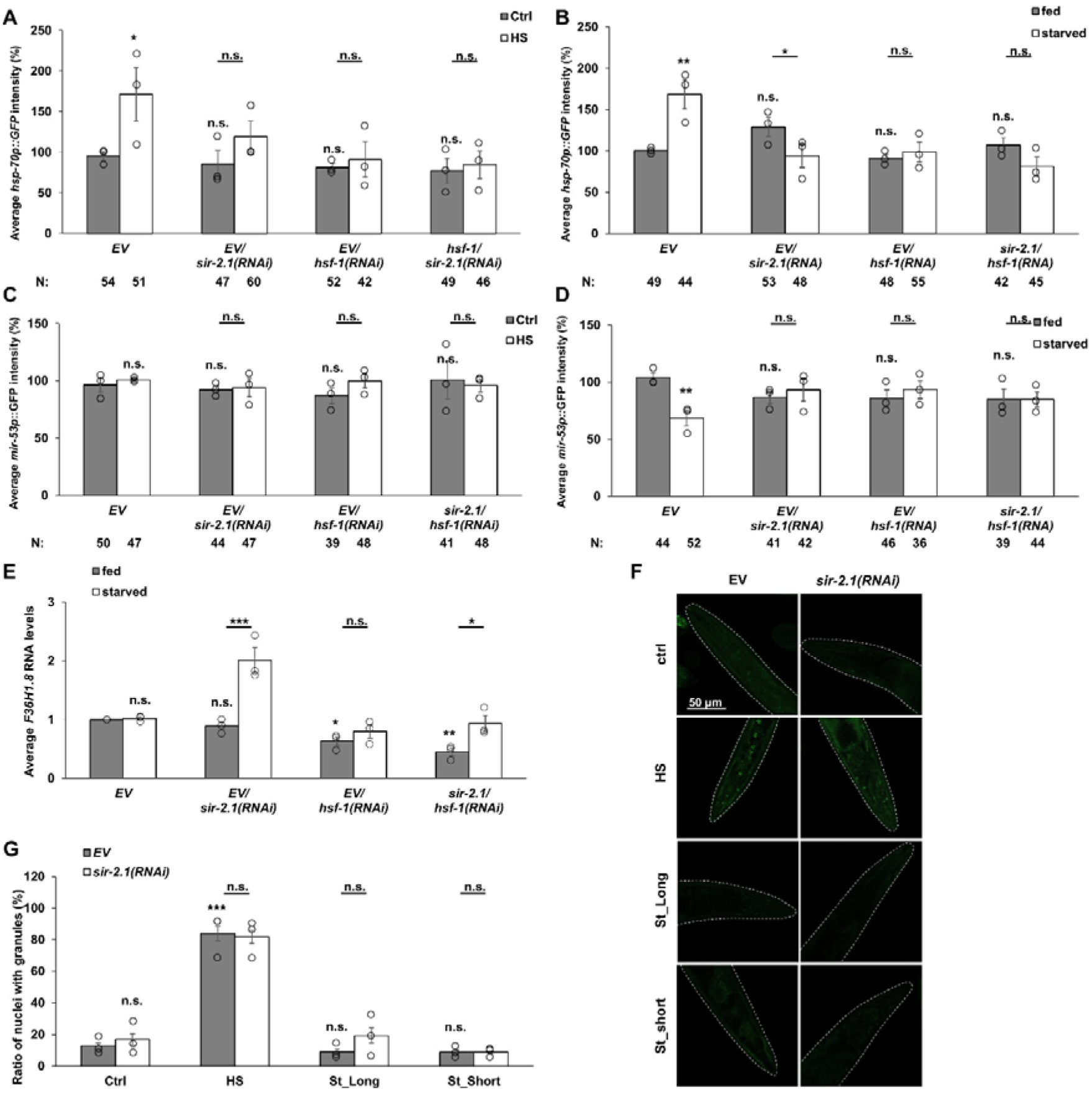
A context-dependent interaction of SIR-2.1 and HSF-1 regulates *miR-53* expression during starvation. (A-B) *hsp-70* promoter activation in response to heat-shock (HS) and starvation upon *sir-2.1* and/or *hsf-1* silencing. (C-D) *mir-53* promoter activation in response to heat-shock and starvation upon *sir-2.1* and/or *hsf-1* silencing. (E) *mir-53* pre- miRNA (*F36H1.8*) levels upon starvation and *sir-2.1* and/or *hsf-1(RNAi)* treatment. (F-G) Representative fluorescence microscopy images (F) and quantification of (G) stress granule formation of HSF-1::GFP in response to 1-hr 35°C heat-shock (HS), 18-hr long-term (ST_Long) and a 2-hr short-term (St_Short) starvation upon EV or *sir-2.1(RNAi)*. Data are expressed as mean ± SEM of three independent experiments. Circles represent the mean values of individual experiments. N, number of animals measured/condition. p values were obtained by two-way ANOVA using the Fisher’s LSD test. n.s.: not significant; *: p<0.05; **: p<0.01; ***: p<0.001.

It was reported that HSF-1 forms nuclear stress granules in response to proteotoxic stresses such as the archetypal heat shock ^53^. We hypothesized that the redistribution of HSF-1 by SIR-2.1 into stress granules might interfere with its availability to promote *miR-53* expression. Therefore, utilizing a strain that expresses HSF-1 as a fusion protein bound to GFP, we monitored the subnuclear distribution of HSF-1 during a 1-hr HS at 35°C, a usually employed 18-hr and a short, 2-hr starvation to prevent a potential re-distribution. In accordance with the data of the Lamitina group ^53^, heat shock led to a significant increase in the number of HSF-1::GFP in stress granules, while neither starvation protocols did so (Figure 5F, G). Moreover, the recruitment of HSF-1 into stress granules was entirely independent of *sir-2.1*. Altogether, these results show that the interaction of HSF-1 and SIR-2.1 and the differential expression of *hsp-70* and *miR-53* during starvation occurs independently of stress granule formation. Thus, the mechanism underlying the context- dependent SIR-2.1/HSF-1 interaction resulting in the downregulation of *mir-53* is yet unclear.

### The Sir2 and PKA pathways converge on ATGL-1 for effective lipolysis

The cAMP-dependent protein kinase regulates lipolysis in response to systemic adrenergic signals^54^. The *C. elegans* catalytic subunit ortholog *kin-1* facilitates lipid mobilization through phosphorylation of ATGL-1^16^ and its lipolytic activity is required for survival under low temperatures and for longevity^17^. Hence, we set out to investigate potential links between the SIR-2.1 and the KIN-1 pathways. First, we stained lipid stores in fed and starved *kin-*1 mutants and confirmed that *kin-1* was also required for lipolysis (Figure 6A). The complete inhibition of lipolysis in the absence of *sir-2.1* or *kin-1* suggests that Sir2 and PKA pathways are equally indispensable and cooperate in lipolysis regulation in the wildtype. Loss of the PKA regulatory subunit KIN-2 results in a constitutive, excessive activation of KIN-1^55^ ^16^. In agreement with this, *kin-2* mutation phenocopied starvation-induced lipid store depletion in *ad libitum* fed worms (Figure 6A). This effect was not influenced by *sir-2.1* or *hsf-1* knockdown (Figures 6A-C), which suggests that KIN-1 acts downstream from the SIR-2.1 pathway.

**Figure 6.**
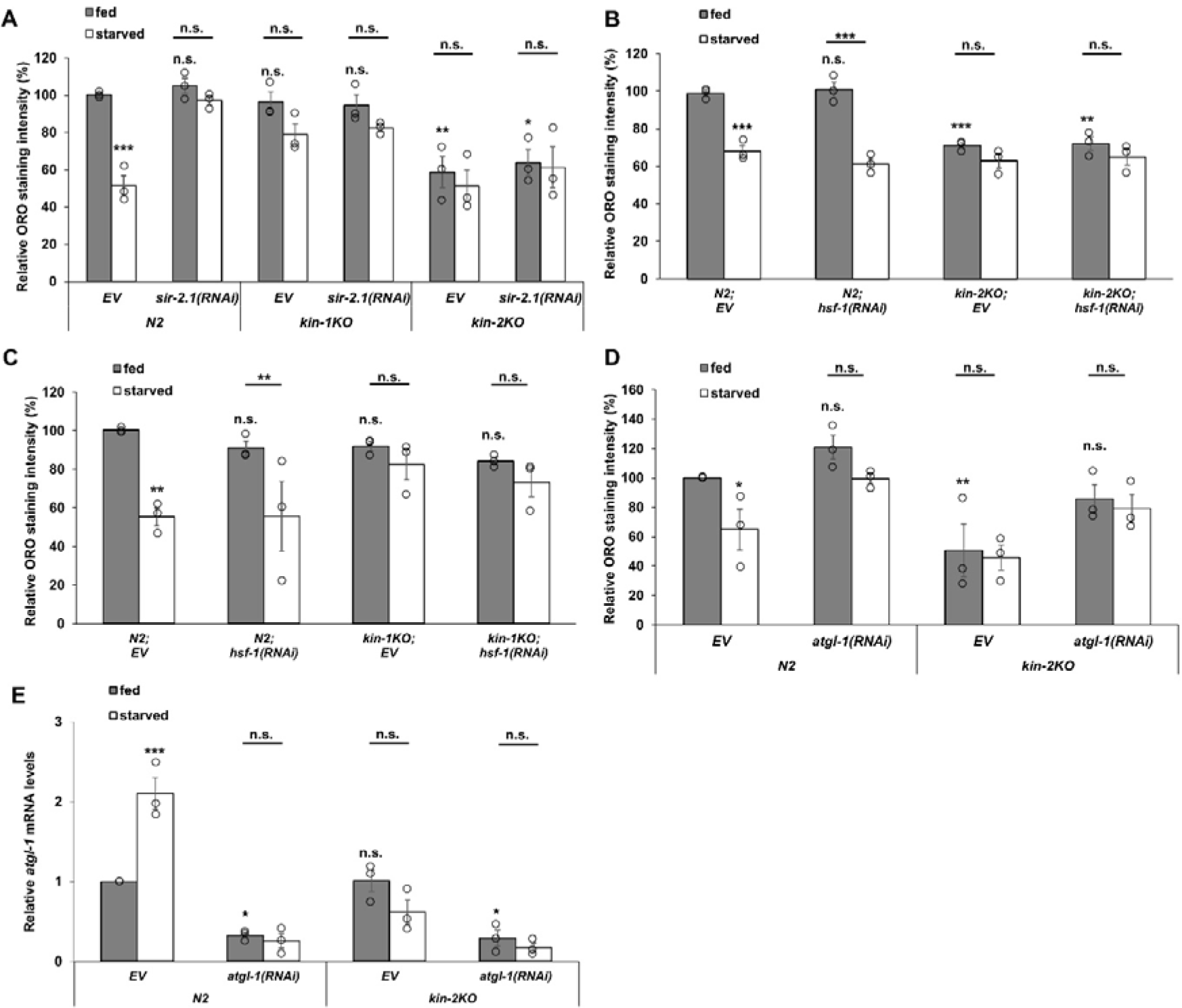
The protein kinase A KIN-1 affects lipolysis and ATGL-1 function downstream from SIR-2.1 and HSF-1. (A) Effect of *kin-1* and *kin-2* mutations on starvation-induced lipid mobilization in EV and *sir-2.1(RNAi)* fed animals. (B) Starvation- induced lipid mobilization in wildtype and *kin-2* mutant animals fed with EV or *hsf- 1(RNAi)*. (C) Starvation-induced lipid mobilization in wildtype and *kin-1* mutant animals fed with EV or *hsf-1(RNAi)* bacteria. (D) Starvation-induced lipid mobilization in wildtype and *kin-2* mutant animals fed with EV or *atgl-1(RNAi)* from hatching. (E) *atgl-1* mRNA levels in wildtype and kin-2 mutant animals fed with EV or *atgl-1(RNAi)* bacteria. Data are expressed as mean ± SEM of three independent experiments. Circles represent the mean values of individual experiments. p values were obtained by two-way ANOVA using the Fisher’s LSD test. n.s.: not significant; *: p<0.05; **: p<0.01; ***: p<0.001.

Next, we re-examined whether KIN-1 uses ATGL-1 to mobilize lipid stores. We employed RNAi against *atgl-1* and measured ORO staining in *kin-2* mutant animals in response to starvation. When the treatment was applied from L4 stage, as in all previous experiments in order to avoid any potential developmental effects, *atgl-1(RNAi)* raised the lipid content in *kin-2* mutants, although not reaching the level of the wildtype (Figure S4). In *kin-2* mutants, KIN-1 exhibits a high activity from fertilization, therefore we employed *atgl-1* RNAi from hatching. Indeed, this treatment resulted in an ORO staining intensity of animals that was no longer significantly different from wildtype (Figure 6D). These results indicate that both SIR-2.1 and KIN-1 converge on ATGL-1. We reasoned that if KIN-1 acts downstream of ATGL-1 post-transcriptional regulation, its activation will not upregulate *atgl-1* mRNA upon starvation. Indeed, the induction was not observed neither in fed, nor in starved *kin-2* mutant animals (Figure 6E), excluding the transcriptional and post-transcriptional regulation of *atgl-1*. Consistent with the kinase activity of KIN-1 and with the reported phosphorylation and stabilization of ATGL-1 protein ^16^, our results confirm that it affects ATGL-1 activity at the post-translational level. We also hypothesize that there may be a negative feedback-loop, which limits *atgl-1* expression when lipase activity and/or lipolysis is sufficient.

### Intestinal proteostasis disturbance inhibits lipid mobilization via *hsf-1*

Given the role HSF-1 plays in proteostasis maintenance, we asked how proteotoxic stress affects lipid mobilization. To this end, before starvation we pre-exposed animals to heat shock, which rapidly induces global protein misfolding and HSF-1 activation^53,56^. The results of ORO staining experiments demonstrated that heat shock completely prevented the starvation-induced drop in lipid stores (Figure 7A). We also monitored the expression of the ATGL-1::GFP reporter and found that heat shock pretreatment inhibited its activation, which was dependent on *hsf-1* (Figure 7B). To establish more specifically whether this phenomenon is indeed caused by misfolded proteins or by other heat shock induced processes, we employed transgenic nematode strains, that constitutively express single misfolded proteins in the body wall muscle or intestine and determined their lipid stores in response to starvation. Animals expressing the muscle-specific amyloid β were reported to be temperature-sensitive with adult onset paralysis and egg-laying deficiency when raised at 20°C^57^. The C-terminal addition of a degron sequence to GFP (GFP::degron) also led to aggregation and paralysis when expressed in muscle^58^. An 82 amino-acid long polyglutamine peptide fused to YFP (Q82::YFP) expressed in the intestine formed aggregates throughout the lifespan^59^. Neither amyloid β nor GFP::degron inhibited significantly the starvation-induced depletion of intestinal lipid stores (Figure 7C and D). In contrast, the intestine-specific expression of Q82::YFP prevented the animals to mobilize their lipid reserves (Figure 7E), which was rescued by *hsf-1(RNAi)* (Figure 7F) suggesting the requirement of HSF-1 (Figure 7F). These findings are in accordance with previously demonstrated tissue-specificity of the SIR-2.1- HSF-1 lipolysis-regulatory axis (Figure 2D and E) and indicate that proteostasis disturbances in the gut inhibit lipolysis and therefore modulate energy metabolism during starvation.

**Figure 7.**
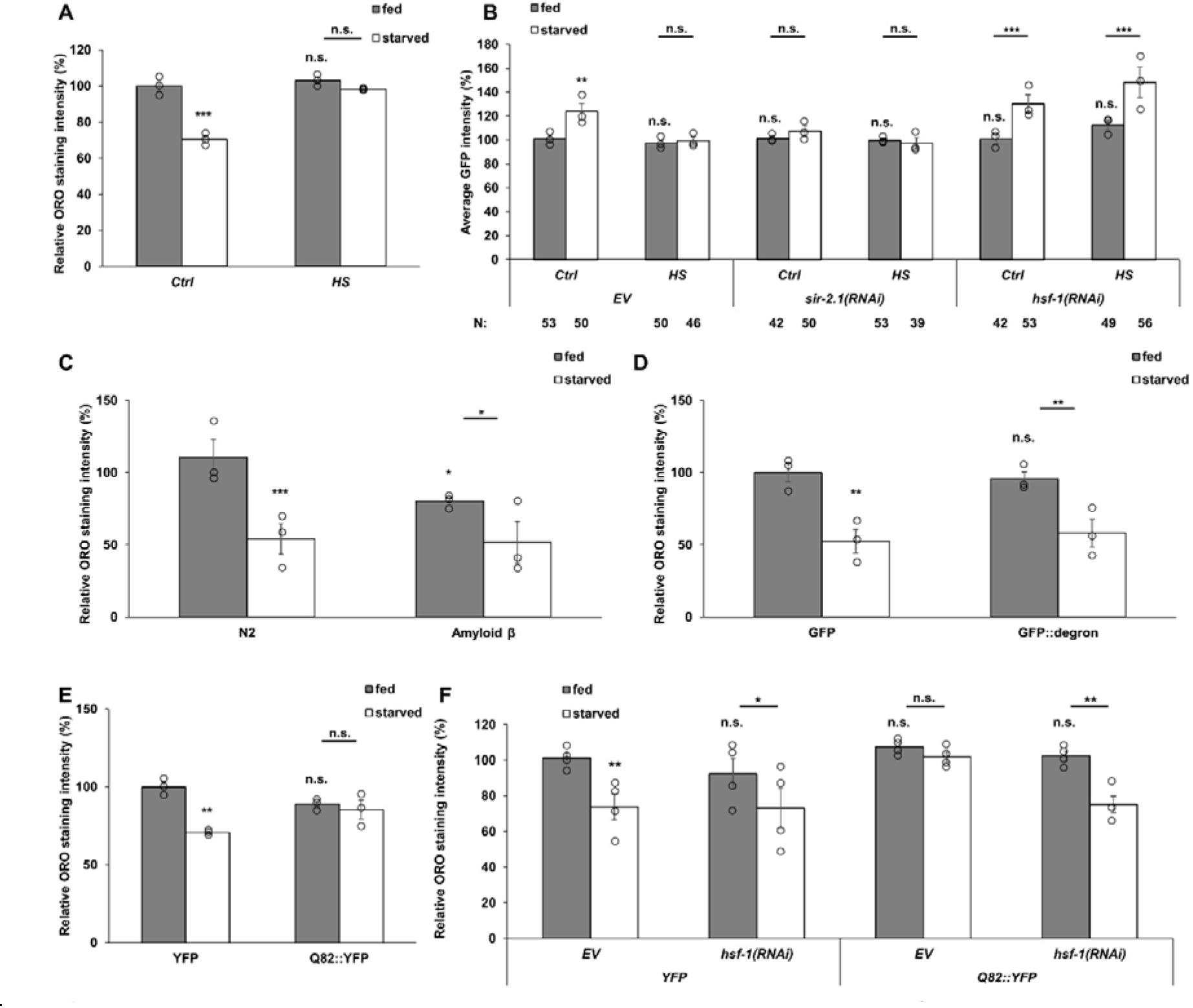
Systemic and intestinal proteotoxic stresses inhibit lipid mobilization during starvation in an HSF-1-dependent manner. (A) The effect of heat shock on lipid stores in response to starvation. (B) The effect of heat shock on the expression of ATGL-1::GFP in response to starvation in worms fed by EV, *sir-2.1* or *hsf-1(RNAi)*. (C-E) The effect of muscle-expressed amyloid ß (C) and GFP::degron (D) and intestinally expressed Q82::YFP (E) misfolded polypeptides on starvation-induced lipid mobilization. (F) The effect of intestinally expressed Q82::YFP on starvation-induced lipid mobilization in worms fed by EV or *hsf-1(RNAi)*. Data are expressed as mean ± SEM of three (A-E) or four (F) independent experiments. Circles represent the mean values of individual experiments. N, number of animals measured/condition. p values were obtained by two-way ANOVA using the Fisher’s LSD test. n.s.: not significant; *: p<0.05; **: p<0.01; ***: p<0.001.

### HSF-1 modulates age-related fat deposition through ATGL-1

HSF-1 has been associated with the aging process. Specifically, loss of HSF-1 promotes, whereas its overexpression retards aging in *C. elegans*^33,39,60^. Further, an age-related decline in HSF-1 activity induced the collapse of proteostasis in nematodes at day 4 of adulthood, which was prevented by HSF-1 overexpression^61^. Moreover, a recent study showed that *hsf-1* RNAi diminished, while HSF-1 overexpression driven by the *sur-5* promoter increased fat deposition in fed 5-day old worms^41^. That finding appeared to support an inhibitory role of HSF-1 on ATGL-1. However, we found that loss of *hsf-1* by gene mutation or silencing, compared to N2, did not reduce lipid stores of day 3 adults in the fed state (see Figures 2, 3 and 6). Hence, we investigated how the aging process affects fat stores and the roles of HSF-1 and ATGL-1. To this end, we monitored the ORO staining intensities of 1, 4 and 8-day old wildtype, *hsf-1* knock-out (*hsf-1KO*) and *hsf-1* overexpressor (*hsf-1OE)* strains driven by the *sur-5* promoter, which were either fed by empty vector (*EV*) or *atgl-1(RNAi)*. Besides the age-related decline in *hsf-1* activity, these age groups were selected keeping in mind a decline in motility after day 3 together with a relatively maintained pharyngeal pumping rate^62,63^, representing decreasing energy expenditure with a constant food consumption, characteristic to early aging.

Consistent with the above, we observed an age-dependent increase in the lipid stores of wild-type worms from day 1 through day 8 of adulthood (Figure 8A-C). *atgl-1*-silenced, compared to EV-fed worms, exhibited similar ORO intensities at days 1 and 4, which became markedly higher by day 8, consistent with the liberation of *atgl-1* from the HSF-1-mediated suppression of feeding worms during aging (Figure 8A-C). The ORO signal of *hsf-1KO*, compared to N2, animals was higher at day 1, which did not show a pronounced increase at later time points (Figure 8D-F). This might be due to pleiotropic effects of HSF-1 on NHR-49^41^, ATGL-1 and a recently discovered requirement of HSF-1 for mitochondrial beta oxidation^64^. Notably, *atgl- 1* silencing elicited a substantial increase in lipid accumulation in *hsf-1KO*, which appeared at day 4 and was more pronounced than that of the wildtype also at day 8 (Figure 8A-F). *hsf- 1OE* nematodes exhibited higher ORO signal than N2 at all ages, which already peaked at day 4 (Figure 8G-I). Importantly, *atgl-1(RNAi)* did not further increase the lipid content in *hsf- 1OE* animals (Figure 8E-F), showing that the genetic activation of *hsf-1* promotes age-related fat deposition via the inhibition of ATGL-1. Altogether these findings reinforce the idea that HSF-1 function is a regulator of lipolysis in adulthood and suggests that its decline limits fat accumulation during aging.

**Figure 8.**
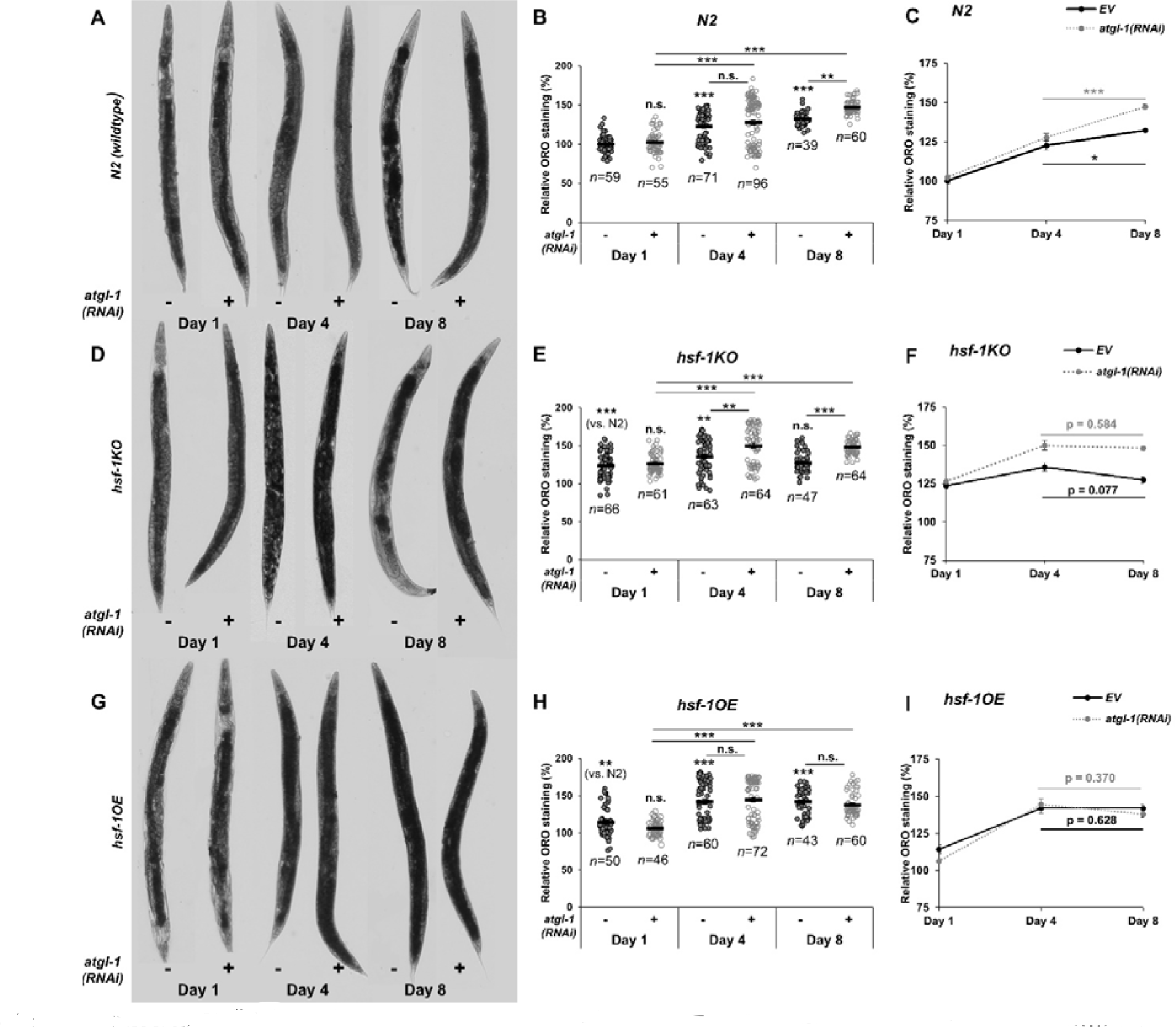
HSF-1 modulates age-related fat deposition through ATGL-1. ORO staining (A, D, G), quantification (B, E, H) and time-dependence (C, F, I) of lipid content in N2 wildtype (A-C), *hsf-1* mutants (*hsf-1KO)* (D-F) and *hsf-1* overexpressors driven by the *sur-5* promoter (*hsf-1OE*) (G-I). Mean ± SEM for a minimum of 39 animals per condition across three replicate experiments. Circles represent individual animals measured. p values were obtained by Kruskal–Wallis test. n.s.: not significant; *: p<0.05; **: p<0.01; ***: p<0.001.

## Discussion

In this study, we demonstrated that regulation of fat mobilization in the *C. elegans* intestine involves the HSF-1 heat-shock transcription factor. Our results show that in response to starvation, the SIR-2.1 deacetylase suspends the HSF-1-mediated repression of adipose triglyceride lipase ATGL-1 expression. This repression occurs at the post-transcriptional level and is conveyed by microRNAs, among which *mir-53* was found to suppress lipolysis (Figure 9). The repression of ATGL-1 by HSF-1 not only inhibits lipolysis during feeding, but also hinders starvation-induced lipolysis under proteostasis disturbance, which promotes fat deposition in adulthood. The decline of HSF-1 function limits fat accumulation through ATGL-1 derepression in aged worms.

**Figure 9.**
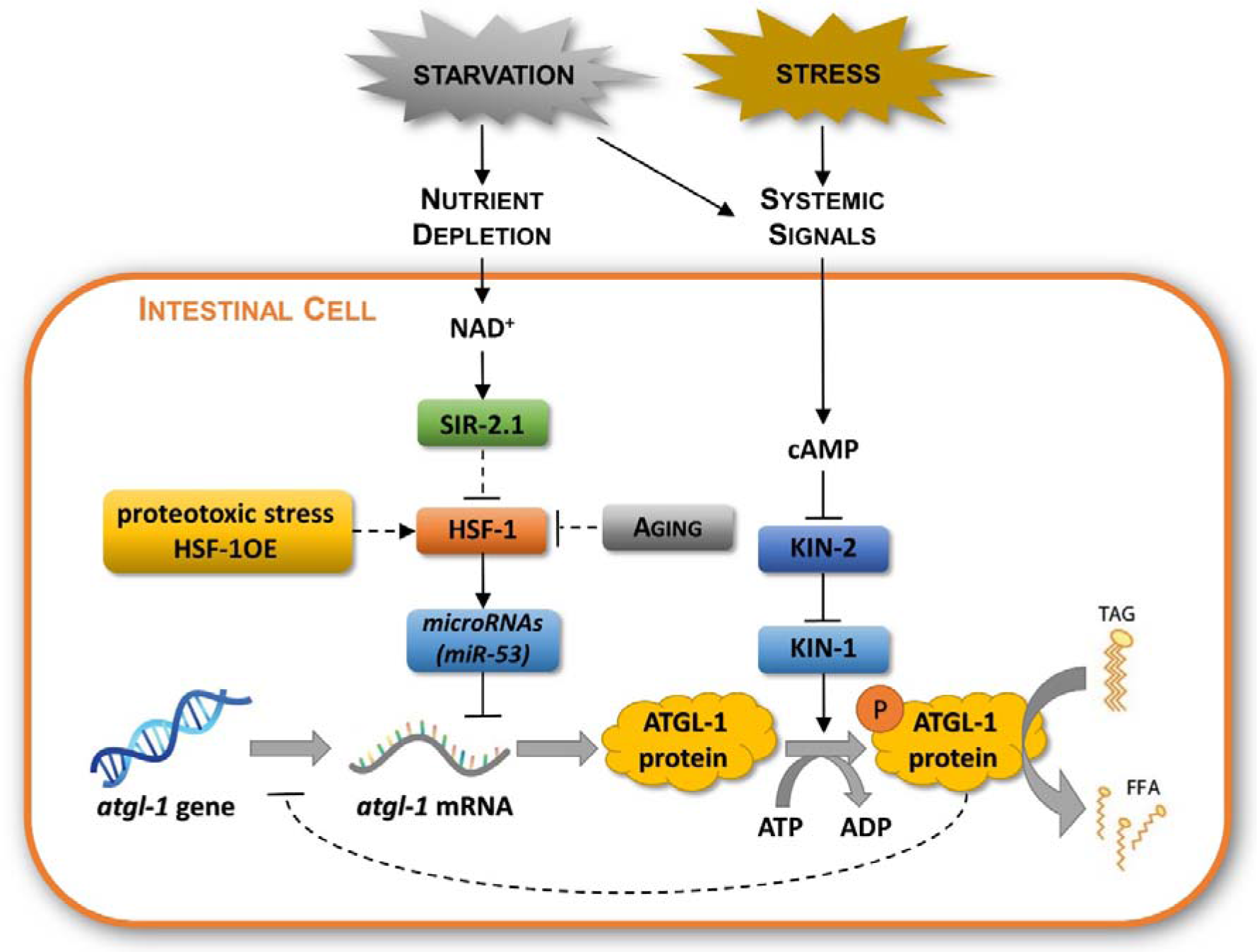
Proposed model of intestinal lipolysis regulation by the Sir2 and PKA pathways converging on ATGL-1. During starvation, nutrient depletion prompts SIR-2.1 to stimulate lipolysis by liberating an HSF-1-dependent, microRNA-mediated suppression of ATGL-1 expression. ATGL-1 protein activity is also regulated by the PKA pathway in response to systemic neuroendocrine signals evoked by starvation and stress. Increased HSF-1 activity by overexpression and by intestinal proteotoxic stress inhibits, its age-related decline promotes lipid mobilization.

In *C. elegans*, energy is stored mostly as lipids: they can make up 35% of the dry body mass of animals, compared to only 3.3% for carbohydrates^65^. 40-55% of total lipid content are made up by triacylglyceride fat stores^66^. This disproportionately large share renders lipid metabolism a critical determinant of energy homeostasis. This role requires a fine-tuned regulation to provide instant energy in stressful conditions and to preserve energy for long-term survival. The fact that developmentally arrested, non-feeding dauer larvae also depend mainly on lipid stores for survival and longevity^67^ provides experimental support for the importance of lipolysis regulation.

Our findings show a direct regulation of lipolysis by HSF-1. Recently, nematode HSF-1 was shown to indirectly affect lipid metabolism through stabilizing the enteric actin network, which promotes membrane trafficking, cellular nutrient absorption and lipid accumulation in the fed state^41^. Another recent study reported the loss of *hsf-1* inhibits mitochondrial β- oxidation and distorts the lipid landscape^64^. These studies together highlight a role for *C. elegans* HSF-1 in ensuring optimal lipid and energy resources in the fed state. This non-canonic role of HSF-1 in lipid regulation appears to be a novel gain of function in metazoa, since in *Saccharomyces cerevisiae*, reduced expression of *hsf1* did not affect lipid droplet abundance^68^.

A crosstalk between HSF-1 and lipids may also be relevant in adaptation to high temperatures, where proteins and membranes are equally subject to changes. Indeed, neuronal HSF-1 promotes survival at high temperatures by extensive fat remodeling in peripheral tissues through a decrease in intestinal fat desaturase and lysosomal lipase expression^69^. Interestingly, HSF-1 is also activated by a lipid, specifically, oleic acid and protects at high temperature and proteotoxicity through histone acetylation and expression of protective genes^64,70^. Thus, HSF-1 utilizes different molecular pathways and connections to lipid homeostasis in fed/starved state and to adapt to high temperature.

According to our results, SIR-2.1 liberates *atgl-1* expression by inhibiting the HSF-1- dependent transcription of *mir-53* pre-RNA. Previous studies showed that SIRT1 and SIR-2.1 activate the HSF-1-dependent expression of heat shock protein genes^30,31^. This was also confirmed in this study by monitoring *hsp-70*-expression upon both heat shock and starvation, which indicates that SIR-2.1 also activates a canonical HSF-1 target without heat shock. The finding that *mir-53* promoter activity changed only upon starvation suggests that the effect of SIR-2.1 on HSF-1 is not simply activation or inhibition. Although the exact mechanism remains unclear, we hypothesize that SIR-2.1-induced changes might redirect HSF-1 between different targets. It is known, that certain interactors/co-regulators of HSF-1, like the E2F/DP complex, are capable of inflicting specific changes on the transcriptional activity of HSF-1, where the expression of subsets of target genes involved in larval development is promoted while others remain unaltered^71^. Mammalian HSF1 deacetylation by SIRT1 prolonged the binding of HSF1 to heat-shock promoter of Hsp70^30^. Hence, the proposed mechanism is similar to how post-translational modifications, such as acetylation or phosphorylation, have a significant effect on the transcriptional activity of p53 towards different targets^72,73^, similarly to its close relative, p73^74^. It is also plausible, that other interactors modulate SIR-2.1/HSF-1- dependent transactivation. Besides its canonical *hsp* targets, HSF-1 was shown to activate a plethora of other genes ^36^, among them several microRNAs^35^. These miRNAs in turn can silence their own targets, thus indirectly involving HSF-1 in the inhibition of certain processes, such as growth, epithelium development or cytoskeleton organization^35^. Due to the extensive nature of HSF-1-regulated microRNA network, it is possible that the apparent incomplete effect of *mir-53* on *atgl-1* mRNA level is a result of the involvement of other, yet unknown miR or miRs. Uncovering these possible regulatory RNAs requires further study.

KIN-1, the *C. elegans* ortholog of human PKA catalytic subunit has been reported to phosphorylate and stabilize ATGL-1 and activate lipolysis^16^. The PKA pathway responds to hormonal signals representing stresses, including starvation in both mammals and nematodes^75–77^. Our results confirm the crucial role of KIN-1 and SIR-2.1 and point out the possibility that while KIN-1/PKA pathway incorporates an organismal, neuroendocrine control into lipolysis regulation, SIR-2.1 responds to cellular changes in nutrient availability in the intestine which is in direct correlation with food availability in the gut lumen. Systemic neuroendocrine signals might activate KIN-1/PKA in stress (danger) conditions, when intestinal cells have sufficient NADH levels, but muscle and neural cells need more fuel for the fight-or-flight response. The KIN-1-mediated post-translational modification on pre-existing ATGL-1 proteins enables worms to react instantly, whereas the SIR-2.1-dependent post-transcriptional activation constitutes a slower, sustained response. Interestingly, both processes act on the stability of ATGL-1 mRNA and protein, respectively. Altogether, our findings outline a complex, synergistic interaction of the two pathways in the regulation of lipolysis.

What physiological consequences might bring about the *hsf-1*-dependent regulation of ATGL-1? Apparently, the reduced flux of lipolysis may spare energy reserves for the organism. This reserve is then available among other vital functions to facilitate protein synthesis and proteostasis, both orchestrated by HSF-1^78^. Moreover, greater lipid resources provide a survival advantage in energy consuming infections^79^. Proteostasis and pathogen response both contribute to fitness and longevity^80,81^. Importantly, HSF-1 is a regulator of proteostasis, immunity and lifespan in the wildtype^33,39,60,82,83^, but also mediates lifespan extension induced by a mutation in *daf-2*^84^, by dietary restriction via bacterial dilution^85^ or complete dietary deprivation^38^. The importance of energy preservation by diminished lipid mobilization is illustrated by the loss of AMPK (*aak-1/2*) in *daf-2* dauer larvae, resulting in rapid consumption of lipid stores and premature depletion following vital organ failure, which was prevented by *atgl-1* RNAi^19,86^. In this regard, *daf-*2-activated HSF-1 might operate in a similar manner to AMPK to preserve lipid resources, although such an assumption remains to be tested.

Our findings on the inhibition of lipolysis by HSF-1 overexpression, *hsp-90* RNAi, heat shock, and expression of intestine-specific misfolded proteins suggest that proteotoxic disturbances counteract the effect of nutrient scarcity on lipid (and energy) mobilization via HSF-1. ATGL-1 activity has been shown to augment mitochondrial oxidation and stress^87^. Thus, the HSF-1-mediated inhibition of ATGL-1 may provide a mechanism to spare resources for proteostasis and prevent the aggravation of proteotoxicity by mitochondrial disturbance in young adults. Such a protective role is also reflected in a different mechanism, whereby epigenetic changes in early life activate HSF-1 to reprogram the lipid landscape and metabolism to protect from proteotoxicity^41,64^. In contrast, the programmed decline of HSF-1 activity during aging^61,88^ might result in ATGL-1 upregulation, excessive lipolysis and mitochondrial stress. Consistent with this assumption, 5-day old *hsf-1(RNAi)*-fed worms exhibited severe fat depletion and transcriptional changes in various lipases, whereas HSF-1 overexpressing worms exhibited increased fat reserves, both longevity and lipid excess required *nhr-49*^41^. However, no fat depletion but an extensive remodeling of lipid metabolism, including compromised β-oxidation, was observed in young adult *hsf-1* silenced polyQ expressing nematodes^41,64^. Our experiments in worms of different ages revealed an age-associated deposition of fat, which was mitigated by the decline and loss, and accelerated by the activation, of *hsf-1,* through *atgl-1.* These studies shed light on the complex regulation of lipid metabolism by HSF-1 in different contexts using different pathways, which requires further studies.

Altogether these findings suggest that HSF-1 coordinates lipid and energy metabolism with proteostasis maintenance, which may contribute to lifespan and healthspan regulation. We note that an analogous lipogenic role was described for mammalian HSF1 through the repression of the AMP-activated protein kinase AMPK^89^, illustrating a functional conservation of the crosstalk between HSF1 and lipid metabolism. HSF1 activity also exhibits age-dependent decline in mammals^90–92^. Whether this and/or analogous HSF1-dependent mechanisms modulate mammalian aging and obesity remains to be investigated.

## Materials and methods

### *C. elegans* strains and maintenance

All strains were obtained from CGC. The list of *Caenorhabditis elegans* strains used in this study can be found in Supplementary Table 1. Animals were kept at 20°C using standard *C. elegans* techniques^93^.

### RNA interference

HT115(DE3) *E. coli* strains producing dsRNA against *atgl-1, hosl-1, ech-4, nhr-49, hlh- 30, pha-4, lips-6* and *pash-1* were all purchased from Dharmacon™ reagents. RNAi strains were kindly provided: *dcr-1* by Gary Ruvkun (Harvard University, USA), *hsp-90*^94^ by Eileen Devaney (University of Glasgow, UK), *sir-2.1* and *hsf-1* by Tibor Vellai (Eötvös Loránd University, Budapest, Hungary) (see Supplementary Table 2). In all experiments empty vector (EV) was used as control. RNAi treatment was performed using standard RNAi feeding method according to^95^: RNAi feeding *E. coli* clones were grown overnight in LB medium containing 100 μg/ml ampicillin. Worms were washed 3 times with M9 buffer and placed on plates containing 1 mM IPTG, 50 μg/ml ampicillin and 6.25 μg/ml tetracyclin seeded with *E. coli* HT115 strains harboring the L4440 empty vector (EV) control and specific RNAi vectors, respectively. If not otherwise stated, RNAi treatments were at the L4 larval stage in order to avoid any developmental effects. Measurements were made after 2 days on RNAi bacteria. To control for proper RNAi dosage in double RNAi treatments, overnight cultures of RNAi bacterial strains were mixed in 1:1 ratio, mixing in the empty vector harbouring strain in single RNAi controls.

### Starvation protocol

Synchronized populations were washed 3 times with M9 buffer and placed either on plates containing bacterial food source, or empty plates for 18 hours.

### Lipid staining assays

Worms were harvested by washing from the plates using PBS-T solution. After 3 washing steps animals were resuspended in 600 μL 40 % isopropanol and rocked at room temperature for 3 minutes. Following a 1 minute 560 g centrifugation of the samples and removal of supernatant 600 μL Oil Red O (ORO) working solution (500 mg ORO in 100 ml isopropanol, freshly mixed with destilled water to a 60 % final concentration) was added to each sample and were rocked at room temperature for 2 hours. After another centrifugation at 560 g, pellets were resuspended in PBS-T and rocked for another 30 minutes. Samples were centrifuged again and animals were placed on microscope slides with a 2% agarose pad and imaged using Olympus DP74 Cooled color camera under an Olympus IX73 microsope. The images taken were turned greyscale, inverted and mean signal intensity inside the outlines of individual animals were measured using ImageJ software.

### Free fatty acid quantification

Free fatty acid content was measured using a Free Fatty Acid Quantification kit (Merck; catalog no. MAK044-1KT). After the respective treatments, synchronized adult worms were resuspended in 5% Triton X-100 solution and homogenized using Kontes™ Pellet Pestle™ Motor and Disposable Pellet Pestles. For complete lysis and triglyceride extraction, the homogenates were sonicated and subjected to two cycles of heating (80°C) and cooling (room temperature). Then worm extracts obtained by centrifugation were used to measure the total free fatty acid content according to the manufacturer’s protocol. Samples were normalized to total worm extract protein using Pierce™ BCA Protein Assay Kit (Thermo Scientific™; catalog no. 23225). Free fatty acid content were determined by measuring absorbance at 570 nm using BMG Labtech CLARIOstar Plus plate reader.

### Fluorescence microscopy

At least 50 worms per condition were placed on a 2% agarose pad, and immobilized by adding 25 mM NaN_3_ in M9 buffer for each single experiment. Pictures were taken by a Olympus IX73 microsope with a Olympus DP74 Cooled color camera using a GFP fluorescent filter. GFP intensity inside the outlines of the individual animals was measured in ImageJ software.

### Confocal microscopy

Animals expressing HSF-1::GFP fusion protein were fixed after their respective treatments using cooled ethanol, which was changed to formaldehyde after a few minutes. Samples were placed to 4°C for one hour and DR nuclear dye was used to stain the nuclei of the cells. Images were acquired with a confocal laser microscope (Zeiss Confocal LSM 710, Carl Zeiss AG, Oberkochen, Germany). (Objective: Plan-Apochromat 63×/1.40 Oil DIC M27. Pinhole: 1.01 AU. Laser wavelength: 488 nm. Detection wavelength: 496–564 nm; 673–757 nm).

### mRNA expression analysis

mRNA from well-fed synchronized population of adult worms was isolated using GeneJET RNA Purification Kit (Thermo Scientific). The mRNA was then transcribed into cDNA by RevertAid™ Premium Reverse Transcriptase (Thermo Scientific). qPCR measurements were performed in an ABI 7300 Real-time PCR machine using Maxima™ SYBR Green/ROX qPCR Master Mix (Thermo Scientific). Primer sequences used in this study can be found in Supplementary Table 3. Relative amounts of mRNAs were determined using the Comparative Cycle Treshold Method for quantitation and normalized to beta-actin mRNA levels. Each experiment was performed three times.

### Statistical analysis

Oil Red O staining and Free Fatty Acid contents were examined by two-way ANOVA with Fisher’s Least Significant Difference (LSD) post hoc test unless stated otherwise in the figure legend. qRT-PCR results were analyzed by two-way ANOVA with Fisher’s LSD post hoc test after normalization to controls. Reporter GFP intensities were analyzed by two-way ANOVA with Fisher’s LSD post hoc test. HSF-1 nuclear granule ratios under different conditions were analyzed by two-way ANOVA with Fisher’s LSD post hoc test. Results were extracted from at least three independent experiments and reproducibility was confirmed. Variables were expressed as mean ± standard error of the mean (SEM). Statistical levels of significance are as follows: *:p<0.05, **:p<0.01, ***:p<0.001.

## Acknowledgements

We thank the *Caenorhabditis* Genetics Center for *C. elegans* strains. RNAi against *dcr-1* was received from Gary Ruvkun (Harvard University, USA) while *hsp-90(RNAi)*^94^ was kindly provided by Eileen Devaney (University of Glasgow, UK). RNAi against *sir-2.1* and *hsf-1* were kind gifts from Tibor Vellai (Eötvös Loránd University, Budapest, Hungary). We’d like to thank Wormbase for collecting and providing data on *C. elegans*. We are grateful to Beatrix Gilányi for technical help, and other members of the Stress Group for discussions. C.S. is thankful for the Merit Prize of the Semmelweis University. This work was funded by grants from the Hungarian Science Foundation (OTKA K116525, K147337) to C.S. and (OTKA PD142838) to M.S., from the Semmelweis University (STIA_18_M/6800313263, STIA-KFI-2020/132257/AOMBT/2020) to C.S. and from the Department of Molecular Biology of the Semmelweis University (Baron Munchausen Program 2023/1) to C.S. The funders had no role in study design, data collection and interpretation, or the decision to submit the work for publication.

## Authors’ Contributions

MS and CS conceived the study. MS and CS designed the experiments. MS, SK, GH and JM performed the experiments. MS, GH and CS analyzed the data. MS and CS provided funding. MS, SK and CS wrote the manuscript. All authors read and approved the manuscript.

## Ethics approval and consent to participate

Not applicable.

## Consent for publication

Not applicable.

## Data availability statement

All data supporting the findings of this study are included in the article and its supplementary information.

## References

1. Mutlu, A. S., Duffy, J. & Wang, M. C. Lipid metabolism and lipid signals in aging and longevity. Dev. Cell 56, 1394–1407 (2021).

2. Grabner, G. F., Xie, H., Schweiger, M. & Zechner, R. Lipolysis: cellular mechanisms for lipid mobilization from fat stores. Nat. Metab. 3, 1445–1465 (2021).

3. Fontana, L., Partridge, L. & Longo, V. D. Extending healthy life span-from yeast to humans. Science (80-.). 328, 321–326 (2010).

4. Colman, R. J. et al. Caloric restriction reduces age-related and all-cause mortality in rhesus monkeys. Nat. Commun. 5, 1–5 (2014).

5. Mattison, J. A. et al. Caloric restriction improves health and survival of rhesus monkeys. Nat. Commun. 8, 14063 (2017).

6. Gabel, K. et al. Effects of 8-hour time restricted feeding on body weight and metabolic disease risk factors in obese adults: A pilot study. Nutr. Heal. Aging 4, 345–353 (2018).

7. Dai, Y., Tang, H. & Pang, S. The Crucial Roles of Phospholipids in Aging and Lifespan Regulation. Front. Physiol. 12, 1–7 (2021).

8. Johnson, A. A. & Stolzing, A. The role of lipid metabolism in aging, lifespan regulation, and age-related disease. Aging Cell 18, 1–26 (2019).

9. Ashrafi, K. et al. Genome-wide RNAi analysis of Caenorhabditis elegans fat regulatory genes. Nature 421, 268–272 (2003).

10. Hars, E. S. et al. Autophagy regulates ageing in C. elegans. Autophagy 3, 93–95 (2007).

11. Zaarur, N. et al. ATGL-1 mediates the effect of dietary restriction and the insulin/IGF- 1 signaling pathway on longevity in C. elegans. Mol. Metab. 27, 75–82 (2019).

12. Wang, M. C., O’Rourke, E. J. & Ruvkun, G. Fat metabolism links germline stem cells and longevity in C. elegans. Science (80-.). 322, 957–960 (2008).

13. Folick, A. et al. Lysosomal signaling molecules regulate longevity in Caenorhabditis elegans. Science (80-.). 347, 83–86 (2015).

14. Ramachandran, P. V. et al. Lysosomal Signaling Promotes Longevity by Adjusting Mitochondrial Activity. Dev. Cell 48, 685–696.e5 (2019).

15. Schreiber, R., Xie, H. & Schweiger, M. Of mice and men: The physiological role of adipose triglyceride lipase (ATGL). Biochim. Biophys. Acta - Mol. Cell Biol. Lipids 1864, 880–899 (2019).

16. Lee, J. H. J. H. et al. Lipid Droplet Protein LID-1 Mediates ATGL-1-Dependent Lipolysis during Fasting in Caenorhabditis elegans. Mol. Cell. Biol. 34, 4165–4176 (2014).

17. Liu, F., Xiao, Y., Ji, X.-L., Zhang, K.-Q. & Zou, C.-G. The cAMP-PKA pathway- mediated fat mobilization is required for cold tolerance in C. elegans. Sci. Rep. 7, 1–10 (2017).

18. Xie, M. & Roy, R. AMP-activated kinase regulates lipid droplet localization and stability of adipose triglyceride lipase in C. elegans dauer larvae. PLoS One 10, 1–17 (2015).

19. Narbonne, P. & Roy, R. Caenorhabditis elegans dauers need LKB1/AMPK to ration lipid reserves and ensure long-term survival. Nature 457, 210–214 (2009).

20. Simmons, G. E., Pruitt, W. M. & Pruitt, K. Diverse roles of SIRT1 in cancer biology and lipid metabolism. Int. J. Mol. Sci. 16, 950–965 (2015).

21. Li, X. SIRT1 and energy metabolism SIRT1 is a Cellular Metabolic Sensor. Acta Biochim Biophys Sin 45, 51–60 (2013).

22. Yao, Y., Liu, L., Guo, G., Zeng, Y. & Ji, J. S. Interaction of Sirtuin 1 (SIRT1) candidate longevity gene and particulate matter (PM2.5) on all-cause mortality: a longitudinal cohort study in China. Environ. Heal. A Glob. Access Sci. Source 20, 1–12 (2021).

23. Herranz, D. et al. Sirt1 improves healthy ageing and protects from metabolic syndrome-associated cancer. Nat. Commun. 1, 1–8 (2010).

24. Satoh, A. et al. Sirt1 extends life span and delays aging in mice through the regulation of Nk2 Homeobox 1 in the DMH and LH. Cell Metab. 18, 416–430 (2013).

25. Chen, J. et al. Sirtuins: Key players in obesity-associated adipose tissue remodeling. Front. Immunol. 13, 1–16 (2022).

26. Chakrabarti, P. et al. SIRT1 controls lipolysis in adipocytes via FOXO1-mediated expression of ATGL. J. Lipid Res. 52, 1693–1701 (2011).

27. Burnett, C. et al. Absence of effects of Sir2 overexpression on lifespan in C. elegans and Drosophila. Nature 477, 482–485 (2011).

28. Tissenbaum, H. A. & Guarente, L. Increased dosage of a sir-2 gene extends lifespan in Caenorhabditis elegans. Nature 410, 227–230 (2001).

29. Berdichevsky, A., Viswanathan, M., Horvitz, H. R. & Guarente, L. C. elegans SIR-2.1 interacts with 14-3-3 proteins to activate DAF-16 and extend life span. Cell 125, 1165– 77 (2006).

30. Westerheide, S. D. et al. Stress-inducible regulation of heat shock factor 1 by the deacetylase SIRT. Science (80-.). 323, 1063–1066 (2009).

31. Raynes, R., Leckey, B. D., Nguyen, K. & Westerheide, S. D. Heat shock and caloric restriction have a synergistic effect on the heat shock response in a sir2.1-dependent manner in Caenorhabditis elegans. J. Biol. Chem. 287, 29045–29053 (2012).

32. Walker, A. K. et al. Conserved role of SIRT1 orthologs in fasting-dependent inhibition of the lipid/cholesterol regulator SREBP. Genes Dev. 24, 1403–1417 (2010).

33. Morley, J. F. & Morimoto, R. I. Regulation of Longevity in Caenorhabditis elegans by Heat Shock Factor and Molecular Chaperones. Mol. Biol. Cell 15, 657–664 (2004).

34. Joutsen, J. & Sistonen, L. Tailoring of proteostasis networks with heat shock factors. Cold Spring Harb. Perspect. Biol. 11, a034066 (2019).

35. Brunquell, J., Snyder, A., Cheng, F. & Westerheide, S. D. HSF-1 is a regulator of miRNA expression in Caenorhabditis elegans. PLoS One 12, 1–24 (2017).

36. Brunquell, J., Morris, S., Lu, Y., Cheng, F. & Westerheide, S. D. The genome-wide role of HSF-1 in the regulation of gene expression in Caenorhabditis elegans. BMC Genomics 17, 1–18 (2016).

37. Li, J., Labbadia, J. & Morimoto, R. I. Rethinking HSF1 in Stress, Development, and Organismal Health. Trends Cell Biol. 27, 895–905 (2017).

38. Steinkraus, K. A. et al. Dietary restriction suppresses proteotoxicity and enhances longevity by an hsf-1-dependent mechanism in Caenorhabditis elegans. Aging Cell 7, 394–404 (2008).

39. Hsu, A.-L. L., Murphy, C. T. & Kenyon, C. Regulation of aging and age-related disease by DAF-16 and heat-shock factor. Science 300, 1142–5 (2003).

40. Seo, K. et al. Heat shock factor 1 mediates the longevity conferred by inhibition of TOR and insulin/IGF-1 signaling pathways in C. elegans. Aging Cell 12, 1073–1081 (2013).

41. Watterson, A. et al. Loss of heat shock factor initiates intracellular lipid surveillance by actin destabilization. Cell Rep. 41, 111493 (2022).

42. Elle, I. C. et al. Tissue- and paralogue-specific functions of acyl-CoA-binding proteins in lipid metabolism in Caenorhabditis elegans. Biochem. J. 437, 231–241 (2011).

43. Jo, H., Shim, J., Lee, J. H., Lee, J. & Kim, J. B. IRE-1 and HSP-4 Contribute to Energy Homeostasis via Fasting-Induced Lipases in C. elegans. Cell Metab. 9, 440–448 (2009).

44. Van Gilst, M. R., Hadjivassiliou, H., Jolly, A. & Yamamoto, K. R. Nuclear hormone receptor NHR-49 controls fat consumption and fatty acid composition in C. elegans. PLoS Biol. 3, 0301–0312 (2005).

45. O’Rourke, E. J. & Ruvkun, G. MXL-3 and HLH-30 transcriptionally link lipolysis and autophagy to nutrient availability. Nat. Cell Biol. 15, 668–676 (2013).

46. Lapierre, L. R., Gelino, S., Meléndez, A. & Hansen, M. Autophagy and lipid metabolism coordinately modulate life span in germline-less C. elegans. Curr. Biol. 21, 1507–1514 (2011).

47. Melo, J. A. & Ruvkun, G. Inactivation of conserved C. elegans genes engages pathogen- and xenobiotic-associated defenses. Cell 149, 452–466 (2012).

48. Qadota, H. et al. Establishment of a tissue-specific RNAi system in C. elegans. Gene 400, 166–173 (2007).

49. Kmiecik, S. W. & Mayer, M. P. Molecular mechanisms of heat shock factor 1 regulation. Trends Biochem. Sci. 1–17 (2021) doi:10.1016/j.tibs.2021.10.004.

50. Somogyvári, M., Gecse, E. & Sőti, C. DAF-21/Hsp90 is required for C. elegans longevity by ensuring DAF-16/FOXO isoform A function. Sci. Rep. 8, 12048 (2018).

51. Jan, C. H., Friedman, R. C., Ruby, J. G. & Bartel, D. P. Formation, regulation and evolution of Caenorhabditis elegans 3’UTRs. Nature 469, 97–103 (2011).

52. Brosnan, C. A., Palmer, A. J. & Zuryn, S. Cell-type-specific profiling of loaded miRNAs from Caenorhabditis elegans reveals spatial and temporal flexibility in Argonaute loading. Nat. Commun. 12, 2194 (2021).

53. Morton, E. A. & Lamitina, T. Caenorhabditis elegans HSF-1 is an essential nuclear protein that forms stress granule-like structures following heat shock. Aging Cell 12, 112–20 (2013).

54. London, E., Bloyd, M. & Stratakis, C. A. PKA functions in metabolism and resistance to obesity: Lessons from mouse and human studies. J. Endocrinol. 246, R51–R64 (2020).

55. Lu, X. Y., Gross, R. E., Bagchi, S. & Rubin, C. S. Cloning, structure, and expression of the gene for a novel regulatory subunit of cAMP-dependent protein kinase in Caenorhabditis elegans. J Biol Chem 265, 3293–3303 (1990).

56. Westwood, J., Clos, J. & Wu, C. Stress-induced oligomerization and chromosomal relocalization of heat_shock factor. Nature 353, 822–827 (1991).

57. Link, C. D. Expression of human β-amyloid peptide in transgenic Caenorhabditis elegans. Proc. Natl. Acad. Sci. U. S. A. 92, 9368–9372 (1995).

58. Link, C. D. et al. Conversion of green fluorescent protein into a toxic, aggregation- prone protein by C-terminal addition of a short peptide. J. Biol. Chem. 281, 1808–1816 (2006).

59. Mohri-shiomi, A. & Garsin, D. A. Insulin Signaling and the Heat Shock Response Modulate Protein Homeostasis in the Caenorhabditis elegans Intestine during Infection *. J. Biol. Chem. 283, 194–201 (2008).

60. Garigan, D. et al. Genetic analysis of tissue aging in Caenorhabditis elegans: A role for heat-shock factor and bacterial proliferation. Genetics 161, 1101–1112 (2002).

61. Ben-Zvi, A., Miller, E. A. & Morimoto, R. I. Collapse of proteostasis represents an early molecular event in Caenorhabditis elegans aging. Proc. Natl. Acad. Sci. U. S. A. 106, 14914–14919 (2009).

62. Huang, C., Xiong, C. & Kornfeld, K. Measurements of age-related changes of physiological processes that predict lifespan of Caenorhabditis elegans. Proc. Natl. Acad. Sci. U. S. A. 101, 8084–8089 (2004).

63. Hsu, A. L., Feng, Z., Hsieh, M. Y. & Xu, X. Z. S. Identification by machine vision of the rate of motor activity decline as a lifespan predictor in C. elegans. Neurobiol. Aging 30, 1498–1503 (2009).

64. Oleson, B. J. et al. Early life changes in histone landscape protect against age- associated amyloid toxicities through HSF-1-dependent regulation of lipid metabolism. *Nat*. Aging 4, 48–61 (2023).

65. Cooper, A. F. & Van Gundy, S. D. Metabolism of Glycogen and Neutral Lipids by Aphelenchus avenae and Caenorhabditis sp. in Aerobic, Microaerobic, and Anaerobic Environments. J. Nematol. 2, 305–315 (1970).

66. Ashrafi, K. Mapping out starvation responses. Cell Metab. 3, 235–236 (2006).

67. O’Riordan, V. B. & Burnell, A. M. Intermediary metabolism in the dauer larva of the nematode Caenorhabditis elegans-II. The glyoxylate cycle and fatty-acid oxidation. Comp. Biochem. Physiol. -- Part B Biochem. 95, 125–130 (1990).

68. Breslow, D. K. et al. A comprehensive strategy enabling high-resolution functional analysis of the yeast genome. Nat. Methods 5, 711–718 (2008).

69. Chauve, L., et al. Neuronal HSF-1 coordinates the propagation of fat desaturation across tissues to enable adaptation to high temperatures in C. elegans. PLoS Biology vol. 19 (2021).

70. Zhou, L., Tong, H., Tang, H. & Pang, S. Fatty acid desaturation is essential for C. elegans longevity at high temperature. Mech. Ageing Dev. 200, 111586 (2021).

71. Li, J., Chauve, L., Phelps, G., Brielmann, R. M. & Morimoto, R. I. E2F coregulates an essential HSF developmental program that is distinct from the heat-shock response. Genes Dev. 30, 2062–2075 (2016).

72. Luo, J. et al. Acetylation of p53 augments its site-specific DNA binding both in vitro and in vivo. Proc. Natl. Acad. Sci. U. S. A. 101, 2259–2264 (2004).

73. Oda, K. et al. p53AIP1, a potential mediator of p53-dependent apoptosis, and its regulation by ser-46-phosphorylated p53. Cell 102, 849–862 (2000).

74. Costanzo, A. et al. DNA damage-dependent acetylation of p73 dictates the selective activation of apoptotic target genes. Mol. Cell 9, 175–186 (2002).

75. Palorini, R. et al. Protein Kinase A Activation Promotes Cancer Cell Resistance to Glucose Starvation and Anoikis. PLoS Genet. 12, 1–41 (2016).

76. Zhang, X., Yang, S., Chen, J. & Su, Z. Unraveling the regulation of hepatic gluconeogenesis. Front. Endocrinol. (Lausanne). 10, 1–17 (2019).

77. Sadeghian, F., Castaneda, P. G., Amin, M. R. & Cram, E. J. Functional Insights into Protein Kinase A (PKA) Signaling from C. elegans. Life 12, 1–14 (2022).

78. Clay, K. J., Yang, Y. & Petrascheck, M. Proteostasis is differentially modulated by inhibition of translation initiation or elongation. Elife 12, 1–21 (2023).

79. Dasgupta, M. et al. NHR-49 transcription factor regulates immunometabolic response and survival of caenorhabditis elegans during enterococcus faecalis infection. Infect. Immun. 88, e00130–20 (2020).

80. López-Otín, C., Blasco, M. & Partridge, L. The hallmarks of aging. Cell 153, 1194– 1217 (2013).

81. Kenyon, C. J. The genetics of ageing. Nature 464, 504–12 (2010).

82. Lazaro-Pena, M. I. et al. HSF-1: Guardian of the Proteome Through Integration of Longevity Signals to the Proteostatic Network. *Front*. Aging 3, 1–26 (2022).

83. Singh, V. & Aballay, A. Heat shock and genetic activation of HSF-1 enhance immunity to bacteria. Cell Cycle 5, 2443–2446 (2006).

84. Chiang, W. C., Ching, T. T., Lee, H. C., Mousigian, C. & Hsu, A. L. HSF-1 regulators DDL-1/2 link insulin-like signaling to heat-shock responses and modulation of longevity. Cell 148, 322–334 (2012).

85. Zhang, M. et al. Role of CBP and SATB-1 in aging, dietary restriction, and insulin-like signaling. PLoS Biol. 7, e1000245. (2009).

86. Xie, M. & Roy, R. Increased levels of hydrogen peroxide induce a HIF-1-dependent modification of lipid metabolism in AMPK compromised C. elegans dauer larvae. Cell Metab. 16, 322–335 (2012).

87. Littlejohn, N. K., Seban, N., Liu, C.-C. & Srinivasan, S. A Feedforward Loop Governs The Relationship Between Lipid Metabolism And Longevity. Elife 9, e58815 (2020).

88. Labbadia, J. & Morimoto, R. I. Repression of the Heat Shock Response Is a Programmed Event at the Onset of Reproduction. Mol. Cell 59, 639–650 (2015).

89. Su, K. H., Dai, S., Tang, Z., Xu, M. & Dai, C. Heat Shock Factor 1 Is a Direct Antagonist of AMP-Activated Protein Kinase. Mol. Cell 76, 546–561.e8 (2019).

90. Liu, A. Y., Minetti, C. A., Remeta, D. P., Breslauer, K. J. & Chen, K. Y. HSF1, Aging, and Neurodegeneration. Adv Exp Med Biol. 1409, 23–49 (2023).

91. Kim, H. & Gomez-Pastor, R. HSF1 and Its Role in Huntington’s Disease Pathology. Adv Exp Med Biol. 1410, 35–95 (2023).

92. Trivedi, R. et al. Augmentation of the heat shock axis during exceptional longevity in Ames dwarf mice. GeroScience 43, 1921–1934 (2021).

93. Brenner, S. The genetics of Caenorhabditis elegans. Genetics 77, 71–94 (1974).

94. Gillan, V., Maitland, K., McCormack, G., Him, N. A. I. I. N. & Devaney, E. Functional genomics of hsp-90 in parasitic and free-living nematodes. Int. J. Parasitol. 39, 1071– 81 (2009).

95. Kamath, R. S., Martinez-Campos, M., Zipperlen, P., Fraser, A. G. & Ahringer, J. Effectiveness of specific RNA-mediated interference through ingested double-stranded RNA in Caenorhabditis elegans. Genome Biol. 2, RESEARCH0002 (2001).

