## Supplementary Figures, Methods & Tables 1-3 for "An intestinal Sir2-HSF1-ATGL1 pathway regulates lipolysis in *C. elegans*"

*Supplementary Information*

Milán Somogyvári*, Saba Khatatneh, Gábor Hajdú, Bar Sotil, József Murányi, Csaba Sőti*

*Department of Molecular Biology, Semmelweis University, Budapest, Hungary*

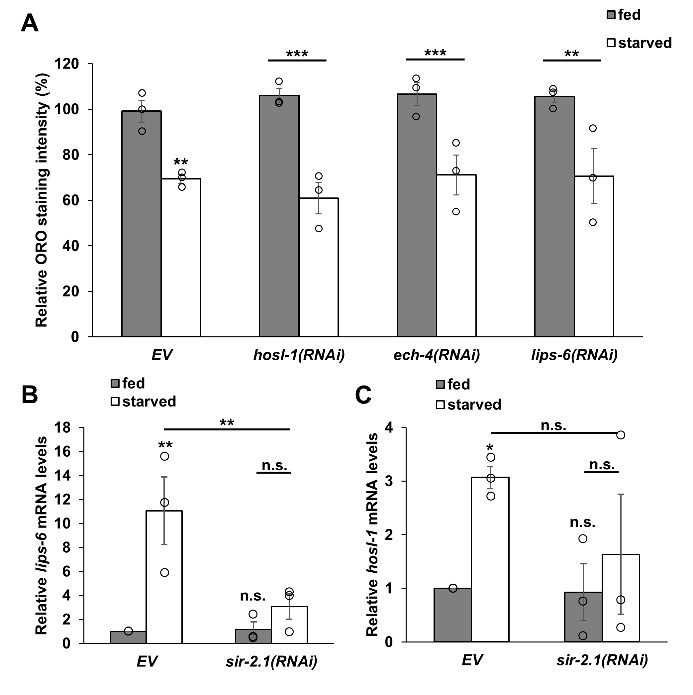
**Supplementary Figures**

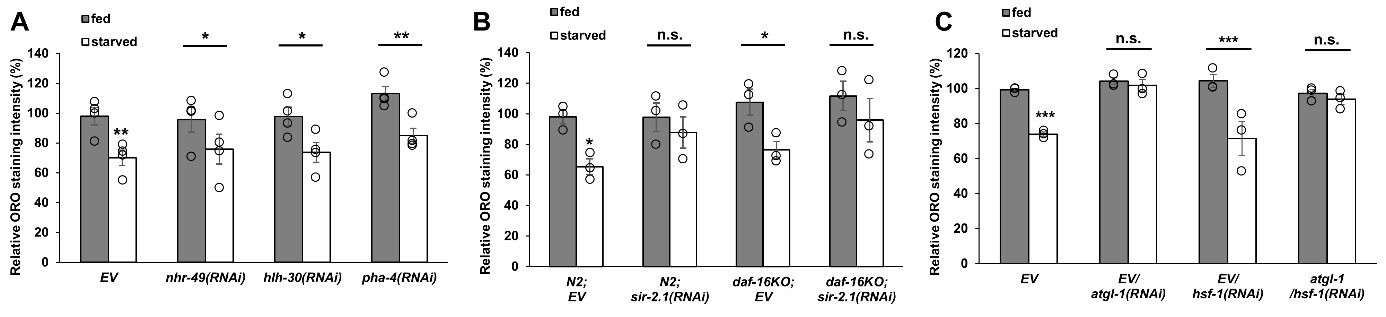
**Figure S1. RNAi knockdown of different triglyceride lipases does not inhibit the reduction of lipid stores, while their induction shows *sir-2.1*-dependency during starvation.** (A) Evaluation of ORO staining of animals treated by RNAi against various lipases. (B-C) Relative mRNA levels of *lips-6* (B) and *hosl-1* (C) on EV and *sir-2.1(RNAi)*. Data are expressed as mean ± SEM. Circles represent the mean values of three independent experiments. p values were obtained by two-way ANOVA using the Fisher’s LSD test. n.s.: not significant; *: p<0.05; **: p<0.01; ***: p<0.001.

**Figure S2. Effect of various transcription factors on starvation-induced lipid mobilization.** (A) RNAi against *nhr-49*, *hlh-30* and *pha-4* did not affect starvation-induced lipid mobilization. (B) Knock-out mutation of *daf-16* did not affect ORO staining, while *sir-2.1(RNAi)* inhibited lipid mobilization in both strains. (C) Lipid mobilization in *atgl-1* and/or *hsf-1* silenced animals. Data are expressed as mean ± SEM. Circles represent the values of three independent experiments. p values were obtained by two-way ANOVA using the Fisher’s LSD test. n.s.: not significant; *: p<0.05; **: p<0.01; ***: p<0.001.

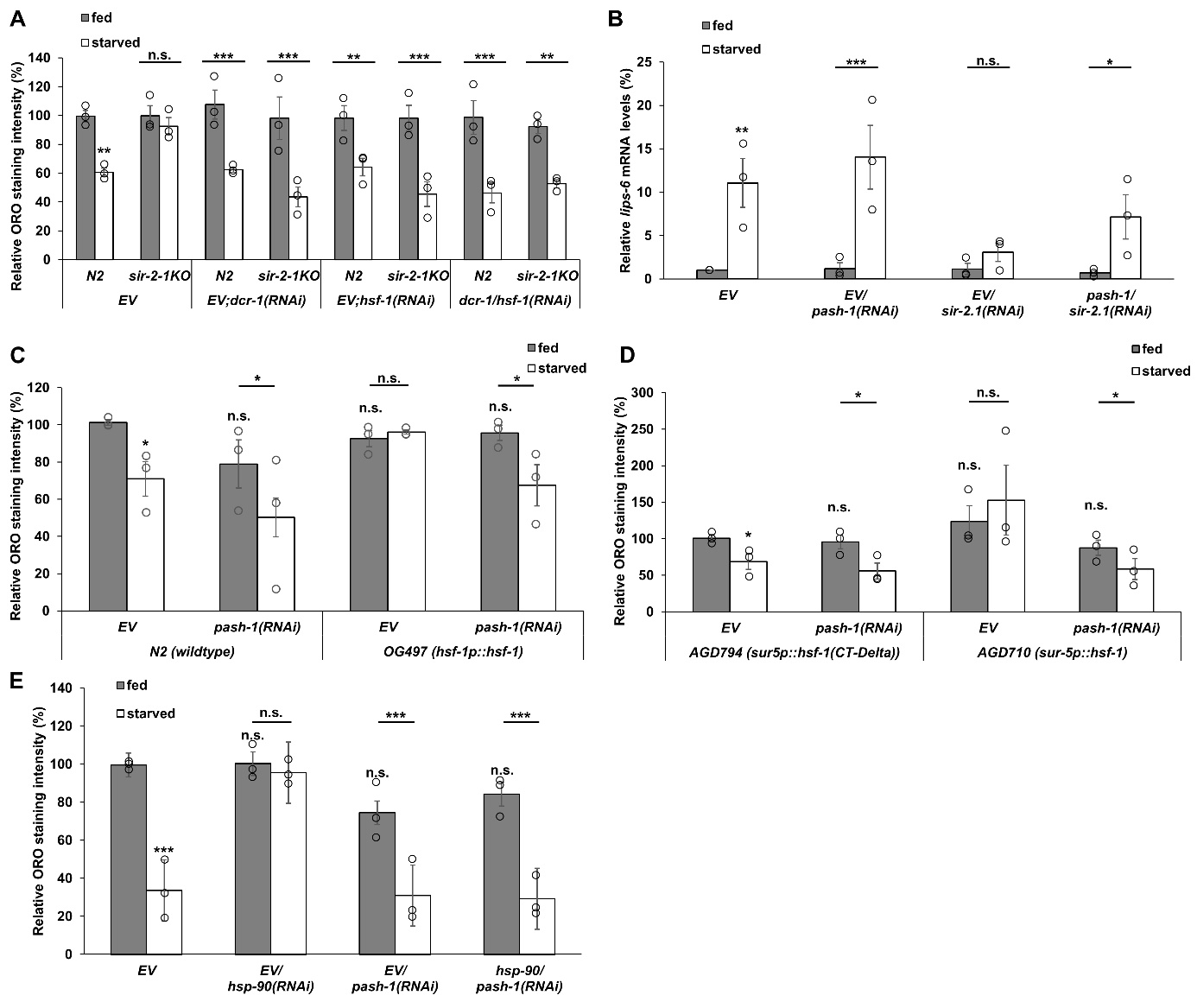
**Figure S3. microRNA formation is required for the HSF-1-dependent inhibition of lipolysis.** (A) The effect of silencing the microRNA maturation gene *dcr-1* on lipid mobilization in wildtype and *sir-2-1* knockout animals. The effect of *pash-1* and/or *sir-2.1* silencing on starvation-induced *lips-6* mRNA levels (B). (C,D) Lipid mobilization in response to starvation in the HSF-1 overexpressor strains upon EV and *pash-1(RNAi). hsf-1* transgene driven by its own promoter is compared to N2 wildtype control (C), whereas *hsf-1* driven by a *sur-5* promoter is compared to its own control expressing the truncated *hsf-1(CT-Delta)* (D). Please note that EV data is identical to that of Figure 3A. (E) Lipid mobilization in response to starvation upon RNAi against *hsp-90* and/or *pash-1*. Data are expressed as mean ± SEM. Circles represent the mean values of three independent experiments. p values were obtained by two-way ANOVA using the Fisher’s LSD test. n.s.: not significant; *: p<0.05; **: p<0.01; ***: p<0.001.

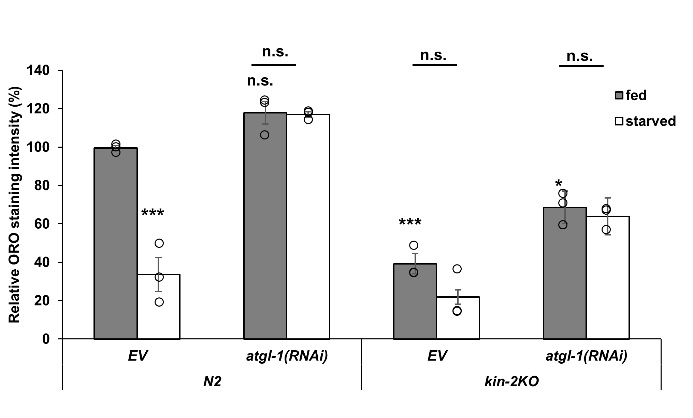
**Figure S4. *atgl-1* RNAi from L4 larval stage partially prevents lipolysis in *kin-2* mutants.** Lipid mobilization in wildtype and *kin-2* mutant animals upon EV or *atgl-1* RNAi treatment from L4 larval stage. Data are expressed as mean ± SEM. Circles represent the mean values of three independent experiments. p values were obtained by two-way ANOVA using the Fisher’s LSD test. n.s.: not significant; *: p<0.05; **: p<0.01; ***: p<0.001.

**Supplementary Tables**

**Supplementary Table 1 - *C. elegans* strains used in this study**

| **Name** | **phenotype** | **genotype** |
| --- | --- | --- |
| N2 | wildtype |  |
| VC199 | sir-2.1KO | *sir-2.1(ok434)* |
| SCS002 | sir-2.1OE | *geIs3 [sir-2.1(+) + rol-6(su1006)]* |
| VS20 | ATGL-1 reporter | *hjIs67 [atgl-1p::atgl-1::gfp + mec-7::RFP]* |
| PS3551 (3 times back-crossed with wildtype) | hsf-1KO | *hsf-1(sy441)* |
| MGH171 | intestinal RNAi sensitive | *sid-1(qt9) V; alxIs9 [vha-6p::sid-1::SL2::GFP]* |
| NR222 | hypodermal RNAi sensitive | *rde-1(ne219) V; kzIs9 [(pKK1260) lin-26p::NLS::GFP + (pKK1253) lin-26p::rde-1 + rol-6(su1006)]* |
| NR350 | muscle RNAi sensitive | *rde-1(ne219) V; kzIs20 [hlh-1p::rde-1 + sur-5p::NLS::GFP]* |
| OG497 | hsf-1OE & reporter | *unc-119(ed3) III; drSi13 [hsf-1p::hsf-1::GFP::unc-54 3'UTR + Cbr-unc-119(+)]* |
| CF1038 | daf-16KO | *daf-16(mu86)* |
| AGD710 | hsf-1OE | *uthIs235 [sur-5p::hsf-1::unc-54 3'UTR + myo-2p::tdTomato::unc-54 3' UTR]* |
| AGD794 | control for AGD710 | *hsf-1 (sy441) I; uthIs225 [sur5p::hsf-1(CT-Delta)::unc-54 3'UTR + myo-2p::tdTomato::unc-54 3' UTR]* |
| MT12989 | mir-53KO | *mir-53(n4113)* |
| MT16471 | mir-60KO | *mir-60(n4947)* |
| MT18037 | mir-75KO | *mir-75(n4472)* |
| VT1605 | mir-53 reporter | *unc-119(ed3) III; maIs234 [mir-53p::GFP + unc-119(+)]* |
| VT1733 | mir-60 reporter | *unc-119(ed3) III; maIs276 [mir-60p::GFP + unc-119(+)]* |
| VL621 | mir-75 reporter | *unc-119(ed3) III; wwIs16 [mir-75p::GFP + unc-119(+)]* |
| BC10060 | hsp-70 reporter | *dpy-5(e907) I; sEx884 [rCesC12C8.1::GFP + pCeh361]* |
| CZ3086 | kin-1KO | *kin-1(ok338)/unc-54(r293)* |
| KG532 | kin-2KO | *kin-2(ce179)* |
| CL2006 | misfolded (amyloid beta) | *dvIs2 [pCL12(unc-54/human Abeta peptide 1-42 minigene) + rol-6(su1006)]* |
| CL2337 | misfolded (degron) | *smg-1(cc546) I; dvIs38 [myo-3p::GFP::degron::3' UTR(long) + rol-6(su1006)]* |
| CL2179 | control for CL2337 | *smg-1(cc546) I; dvIs179 [myo-3p::GFP::3' UTR(long) + rol-6(su1006)]* |
| CB1301 | unc-54ts mutant | *unc-54(e1301)* |
| CB1157 | unc-54ts mutant | *unc-54(e1157)* |
| GF66 | misfolded (polyQ) | *dgEx66 [(pAMS58) vha-6p::Q82::YFP + rol-6(su1006) + pBluescript II]* |
| GF63 | control for GF66 | *dgEx63 [pAMS52 vha-6p::Q0::YFP + rol-6(su1006) + pBluescript II]* |

**Supplementary Table 2 - *RNAi* strains used in this study**

| **Target gene** | **Origin** |
| --- | --- |
| *atgl-1* | Dharmacon™ reagents |
| *hosl-1* |  |
| *ech-4* |  |
| *nhr-49* |  |
| *hlh-30* |  |
| *pha-4* |  |
| *lips-6* |  |
| *pash-1* |  |
| *dcr-1* | Gary Ruvkun (Harvard University, USA) |
| *hsp-90* | Eileen Devaney (University of Glasgow, UK) |
| *sir-2.1* | Tibor Vellai (Eötvös Loránd University, Budapest, Hungary) |
| *hsf-1* |  |

**Supplementary Table 3 - *primers* used in this study**

| **Primer name** | **Sequence** |
| --- | --- |
| *actinFW* | 5'-ATCACCGCTCTTGCCCCATC-3' |
| *actinRV* | 5'-GGCCGGACTCGTCGTATTCTTG-3' |
| *atgl-1FW* | 5’-TCGTTGCCTGTGGTCTCATC-3’ |
| *atgl-1RV* | 5’-ATGCATGGAGCCAATCCACA-3’ |
| *lips-6FW* | 5’-GAAGGAAGGAACCACGAGCA-3’ |
| *lips-6RV* | 5’-TGATGTGACTGCTGTTCGC T-3’ |
| *hosl-1FW* | 5’-AGGCAACTTCAGGACCACTG-3’ |
| *hosl-1RV* | 5’-CTTCCCCTCCATCGGAATCG-3’ |
